# Metric learning enables synthesis of heterogeneous single-cell modalities

**DOI:** 10.1101/834549

**Authors:** Rohit Singh, Brian Hie, Ashwin Narayan, Bonnie Berger

## Abstract

A complete understanding of biological processes requires synthesizing information across heterogeneous modalities, such as age, disease status, or gene/protein expression. Until recently, single-cell profiling experiments could measure only a single modality, leading to analysis focused on integrating information across separate experiments. However, researchers can now measure multiple modalities simultaneously in a single experiment, providing a new data paradigm that enables biological discovery but also requires new conceptual and analytic models. We therefore present Schema, an algorithm that leverages a principled *metric learning* strategy to synthesize multimodal information from the same experiment. To demonstrate the flexibility and power of our approach, we use Schema to infer cell types by integrating gene expression and chromatin accessibility data, perform differential gene expression analysis while accounting for batch effects and developmental age, estimate evolutionary pressure on peptide sequences, and synthesize spliced and unspliced mRNA data to infer cell differentiation. Schema can synthesize arbitrarily many modalities and capture sophisticated relationships between them, is computationally efficient, and provides a valuable conceptual model for exploring and understanding complex biology.

## Introduction

High-throughput assays can now measure diverse cellular properties, including transcriptomic^1–3^, genomic^4,5^, epigenomic^6–8^, proteomic^9^, functional^5^, and spatial^10^ data modalities. Excitingly, single-cell experiments increasingly profile multiple modalities simultaneously within the same experiment^5,6,9,10^, enabling researchers to investigate covariation across modalities; for instance, researchers can study epigenetic gene regulation by correlating gene expression and chromatin accessibility across the same population of cells. Importantly, since the underlying experiments provide us with multimodal readouts, we do not need to integrate modalities across different populations of cells^11–17^.

Simultaneous multimodal experiments present a new analytic challenge of synthesizing agreement and disagreement across modalities. For example, how should one interpret the data if two cells look similar transcriptionally but are different epigenetically? While some multimodal analysis accommodates only specific modalities (e.g., special tools for spatial transcriptomics^18,19,60^), is aimed at just gene-set estimation^20,21^, or is limited only to a pair of modalities^20^, a general multimodal analysis paradigm that applies and extends to any data modality holds the promise of unifying these observations to inform biological discovery. Importantly, given the rapid biotechnological progress that continues to enable novel measurement modalities and easier simultaneous multimodal profiling, such a paradigm should scale to massive single-cell datasets and be robust to noise and sparsity in the data. Furthermore, the ability to synthesize more than just two modalities provides deeper insights and more robust (accurate) inferences, as we demonstrate.

We therefore present Schema, a method that synthesizes multimodal data based on a conceptual framework that accommodates any number of arbitrary modalities. Schema draws from metric learning ^22–25^, the subfield of machine learning concerned with computing an accurate measure of similarity (equivalently, distance) on a dataset. Our critical insight is to interpret each modality as describing a measure of distance among the underlying cells; we can then newly formulate the synthesis problem as reconciling the information implied by these different distance measures. For a modality chosen as the reference, Schema computes a transformation of its data that achieves a desired level of agreement with data from other modalities (**Figure 1**). For example, with RNA-seq data as the reference modality, Schema can transform the data so that it incorporates information from other modalities but remains amenable to standard RNA-seq analyses (e.g. cell-type inference, trajectory analysis, and visualization). A particularly useful property of Schema is that it provides weights that encode the level of agreement between features in the reference modality and the other modalities, enabling biological interpretation of cellular covariation among modalities.

**Figure 1:**
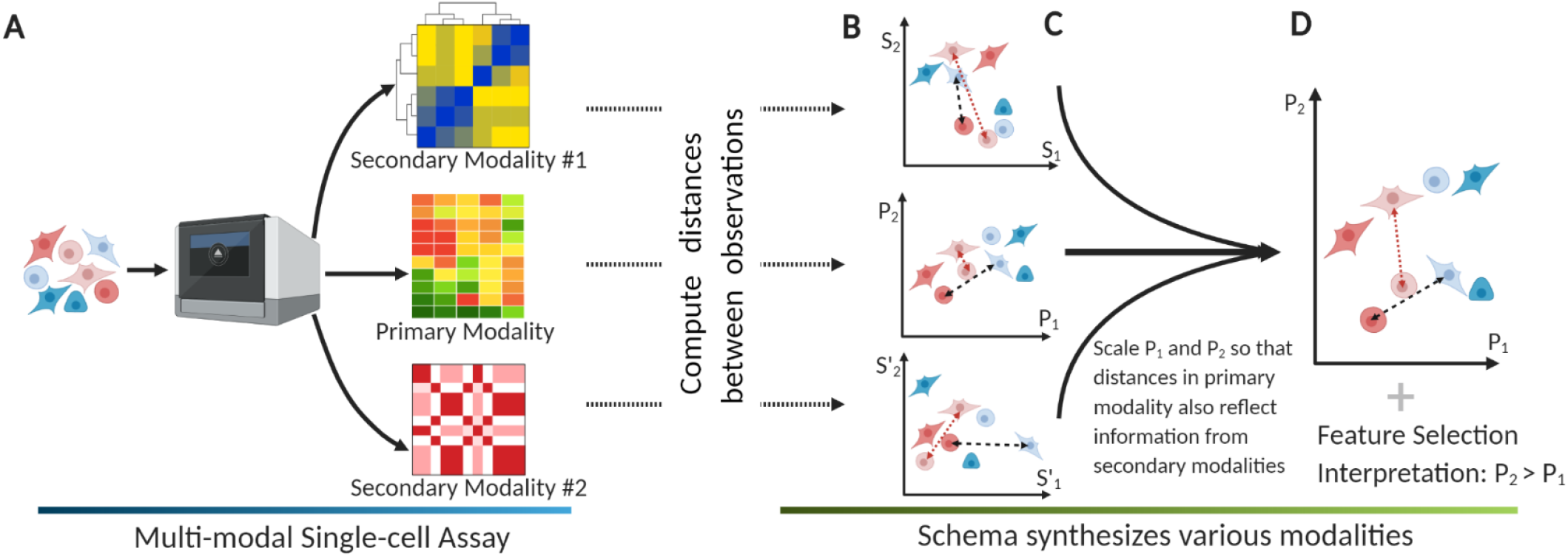
Integration of simultaneously assayed modalities using Schema. (**A**) Schema is designed for assays where multiple modalities are simultaneously measured for each cell. The researcher designates one high-confidence modality as the primary (i.e., reference) and one or more of the remaining modalities as secondary. (**B**) Each modality’s observations are mapped to points in a multi-dimensional space, with an associated distance metric that encapsulates modality-specific similarity between observations. Across the three graphs, the dashed and dotted lines indicate distances between the same pairs of observations. (**C**) Schema transforms the primary-modality space by scaling each dimension so that the distances in the transformed space have a higher (or lower, as desired) correlation with corresponding distances in the secondary modalities; arbitrary distance metrics are allowed for the latter. Importantly, the transformation is provably guaranteed to limit the distortion of the original space, thus ensuring that information in the primary modality is preserved. (**D**) The new point locations represent information synthesized from multiple modalities into a coherent structure. To compute the transformation, Schema weights features in the primary modality by their importance to its objective; we have found this feature-selection aspect very useful in biological interpretation of its results.

In this study, we demonstrate how the generality and utility of Schema can go beyond existing analyses of diverse multimodal datasets and extract novel biological insights. We synthesize RNA-seq and ATAC-seq modalities from multimodal data on 11,296 mouse kidney cells to infer cell types, with Schema enabling an 11% increase in accuracy over previously described approaches. On a dataset of 16,152 embryonic mouse cells from six samples, we use Schema to perform differential expression analysis that accounts for confounding batch effects. By synthesizing spliced and unspliced mRNA counts, we newly infuse RNA “velocity” information into t-SNE embeddings, creating more insightful visualizations of cell differentiation. Schema is designed to support the continually-expanding breadth of single-cell technologies while retaining the power, tunability and interpretability required for effective exploratory analysis.

## Results

### Multimodal synthesis as metric learning

Before the advent of multimodal single-cell experiments, computational analysis has focused on variation within a single modality. A critical problem brought about by the advent of multimodal single-cell experiments is how to reason about information *across* modalities in a mutually consistent way. Our key intuition is that each modality gives us information about the biological similarity among cells in the dataset, which we can mathematically interpret as a modality-specific distance metric. For example, in RNA-seq data, cells are considered biologically similar if their gene expression profiles are shared; this may be proxied as the Euclidean distance between normalized expression vectors, with shorter distances corresponding to greater similarity.

To synthesize these distance metrics, we draw inspiration from *metric learning* (**Supp Text 1**). Given a reference modality, Schema transforms this modality such that its Euclidean distances agree with a set of supplementary distance metrics from the other modalities, while also limiting the distortion from the original reference modality. Analyses on the transformed data will thus incorporate information from *all* modalities (**Figure 1**).

In our approach, the researcher starts by designating one of the modalities as the *primary* (i.e., reference) modality, consisting of observations that are mapped to points in a multi-dimensional space. In the analyses presented here, we typically designate the most informative or high-confidence modality as the primary (i.e., reference), with RNA-seq being a frequent choice (**Discussion**). The coordinates of points in the primary modality are then transformed using information from *secondary* modalities. Importantly, the transformation is constrained to limit the distortion to the primary modality below a researcher-specified threshold. This acts as regularization, preventing Schema from overfitting to other modalities and ensuring that the high-confidence information contained in the primary modality is preserved. We found this constraint to be crucial to successful multimodal syntheses. Without it, an unconstrained alignment of modalities using, for instance, canonical correlation analysis (a common approach in statistics for inferring information from cross-covariance matrices), is prone to overfitting to sample-specific noise, as we show in our results.

To see how Schema’s transformation synthesizes modalities, consider the case where the primary dataset is gene expression data. While the points close in Euclidean space are likely to be biologically similar cells with shared expression profiles, longer Euclidean distances are less informative. Schema’s constrained optimization framework is designed to preserve the information contained in short-range distances, while allowing secondary modalities to enhance the informativity of longer distances by incorporating, for example, cell-type metadata, differences in spatial density, or developmental relationships. To facilitate the representation of complex relationships between modalities, arbitrary distance metrics and kernels are supported for secondary modalities.

Schema’s measure of inter-modality alignment is based on the Pearson correlation of distances, which is optimized via a quadratic programming algorithm, for which further details are provided in **Methods**. An important advantage of Schema’s algorithm is that it returns coefficients that weight features in the primary dataset based on their agreement with the secondary modalities (for example, weighting genes in a primary RNA-seq dataset that best agree with secondary developmental age information). In this study, we demonstrate this interpretability in our applications of Schema.

### Inferring cell types by synthesizing gene expression and chromatin accessibility

We first sought to demonstrate the value of Schema by applying it to the increasingly common and broadly interesting setting in which researchers simultaneously profile the transcriptome and chromatin accessibility of single cells^6^. Focusing on cell type inference, a key analytic step in many single-cell studies, we applied Schema on a dataset of 11,296 mouse kidney cells with simultaneously assayed RNA-seq and ATAC-seq modalities and found that synthesizing the two modalities produces more accurate results than using either modality in isolation (**Figure 2F**, **Supp Figure 3**).

**Figure 2:**
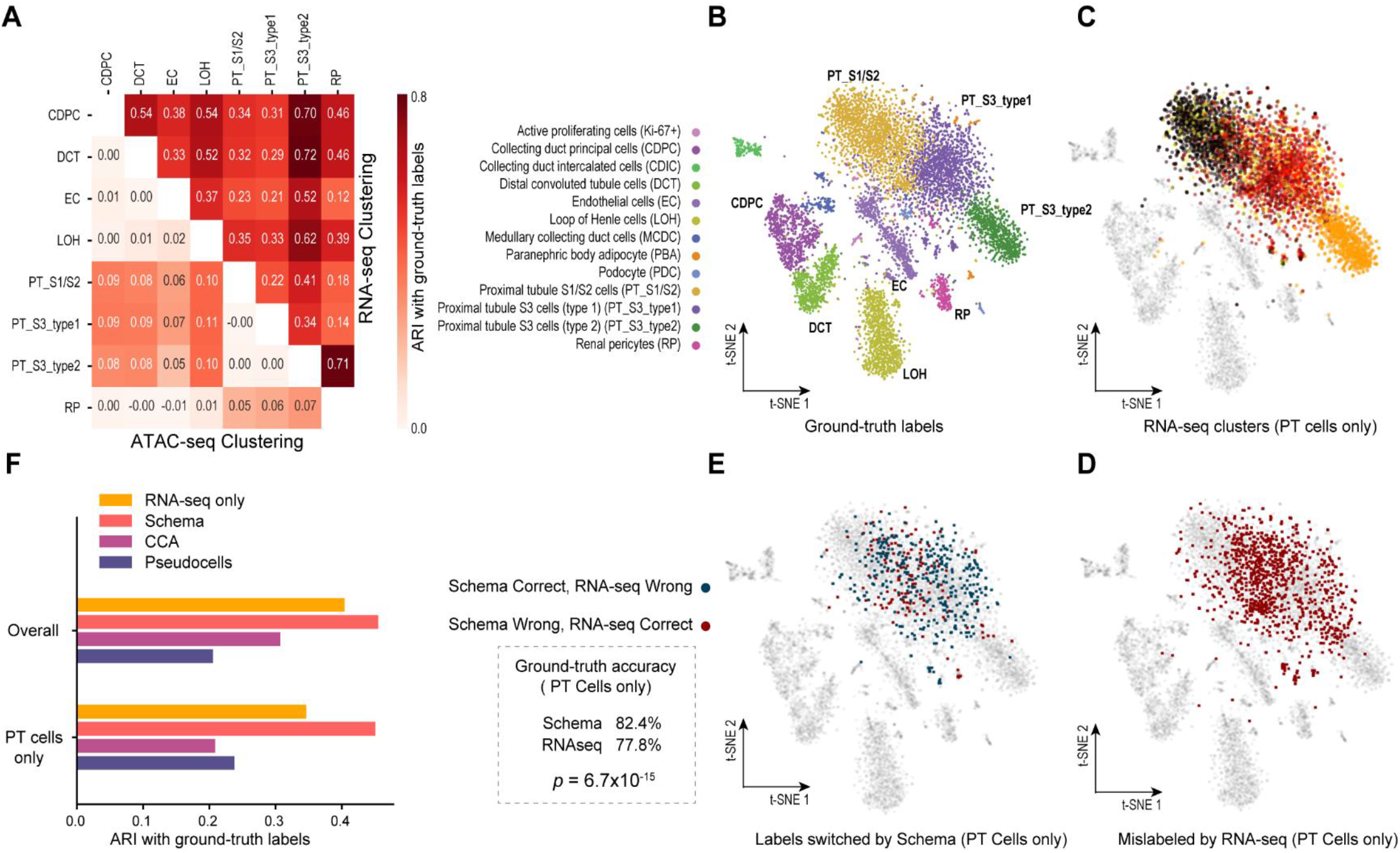
Synthesis of RNA-seq and ATAC-seq information leads to more accurate cell type inference. (**A**) Leiden clustering^50^ of cellular profiles results in greater ARI agreement with ground-truth cell type labels when featurizing cells by RNA-seq profiles alone compared to featurizing with ATAC-seq profiles alone. ATAC-seq does provide relatively more information when distinguishing PT cells. (**B**) Ground truth labels from Cao et al.^6^ (**C, D**) To assess the ground-truth accuracy of Leiden clustering, we assigned each cluster to the cell type most frequently seen in the ground-truth labels of its members. Clusters where labels are more mixed will thus have lower accuracy. Clustering on RNA-seq profiles alone results in many PT cells assigned to such clusters. (**E**) Schema synthesis of RNA- and ATAC-seq features, followed by Leiden clustering, results in significantly greater concordance with ground-truth cell types when compared to Leiden clustering on the RNA-seq features alone (One-sided binomial test, *p* = 6.7 x 10^−15^). (**F**) ARIs of clusters from synthesized data are higher, especially for PT cells. Synthesizing the modalities using canonical correlation analysis (CCA) or a “pseudocell” approach described in the original study (see **Methods**) results in lower ARI scores.

With RNA-seq as the primary (i.e., reference) dataset and ATAC-seq as the secondary, we applied Schema to compute a transformed dataset in which pairwise RNA-seq distances among cells are better aligned with distances in the ATAC-seq peak counts data while retaining a very high correlation with primary RNA-seq distances (≥ 99%, **Methods**). We then clustered the cells by performing Leiden community detection^50^ (a recently introduced modification of the Louvain algorithm^51^) on the transformed dataset and compared these clustering assignments to the Leiden clusters obtained without Schema transformation. We used the expertly defined-labels from Cao et al.^6^ as the ground truth cluster labels and quantified clustering agreement with the adjusted Rand index (ARI), which has a higher value if there is greater agreement between two sets of labels. Leiden clustering on Schema-transformed data better agrees with the ground truth annotations of cell types (ARI of 0.46) than the corresponding Leiden cluster labels using just RNA-seq or ATAC-seq datasets individually (ARIs of 0.40 and 0.04, respectively, **Figure 2F**). Here, Schema facilitated a biologically informative synthesis despite limitations of data quality or sparsity in the ATAC-seq secondary modality. We observed that using only ATAC-seq data to identify cell types leads to poor concordance with ground-truth labels (**Supp Figure 3**), likely because of the sparsity of this modality (for example, only 0.28% of the peaks were reported to have non-zero counts, on average); this sparsity was also noted by the original study authors. We note that an unconstrained synthesis of the modalities using canonical correlation analysis (CCA) resulted in an ARI of 0.31, lower than what is achieved by using just the RNA-seq modality (**Figure 2F**). However, since Schema constrains the ATAC-seq modality’s influence when synthesizing it with RNA-seq data, we could extract additional signal provided by ATAC-seq while preserving the rich information provided by the transcriptomic modality. We also evaluated a heuristic approach described in the original study: group cells into small clusters (“pseudocells”) by RNA-seq similarity and compute an average ATAC-seq profile per pseudocell, using these profiles for the final clustering (**Methods**). This approach also underperformed Schema (ARI of 0.20).

To further analyze why combining modalities improves cell type clustering, we obtained Leiden cluster labels using either the RNA-seq or the ATAC-seq modalities individually. We then evaluated these cluster assignments by iterating over subsets of the data, each set covering only a pair of ground-truth cell types, and used the ARI score to quantify how well the cluster labels distinguished between the two cell types. While RNA-seq clusters have higher ARI scores overall, indicating a greater ability to differentiate cell types, ATAC-seq does display a relative strength in distinguishing proximal tubular (PT) cells from other cell types (**Figure 2A**). PT cells are the most numerous cells in the kidney dataset and many of the misclassifications in the RNA-seq based clustering relate to these cells (**Figure 2B-D**). When the two modalities are synthesized with Schema, a significant number of these PT cells are correctly assigned to their ground truth cell types (one-sided binomial test, *p* = 6.7 x 10^−15^), leading to an overall improvement in clustering quality (**Figure 2E**).

### Differential expression analysis while accounting for batch effects and developmental age

Aside from cell type inference, another important single-cell analysis task that stands to benefit from multimodal synthesis is the identification of differentially expressed marker genes. To illustrate how, we explored a mouse gastrulation single-cell dataset^57^, consisting of 16,152 epiblast cells split over three developmental timepoints (E6.5, E7.0, and E7.25) and with two replicates at each timepoint, resulting in six distinct batches (**Figure 3A**). Applying Schema to this dataset, we sought to identify differentially expressed genes that are consistent with the developmental time course while being robust to batch effects between the replicate pairs. To perform differential expression analysis with Schema, RNA-seq data should be used as the primary modality, while the distance metrics of the secondary modalities specify how cells should be differentiated from each other. Here, we used batch and developmental-age information as secondary modalities, configuring Schema to maximize RNA-seq data’s agreement with developmental age and minimize its agreement with batch information. We weighted these co-objectives equally; results were robust to ± 25% variations in these weights (**Supp Figure 9**). We used RNA-seq data as the primary dataset, representing it by its top ten principal components (**Methods**).

**Figure 3:**
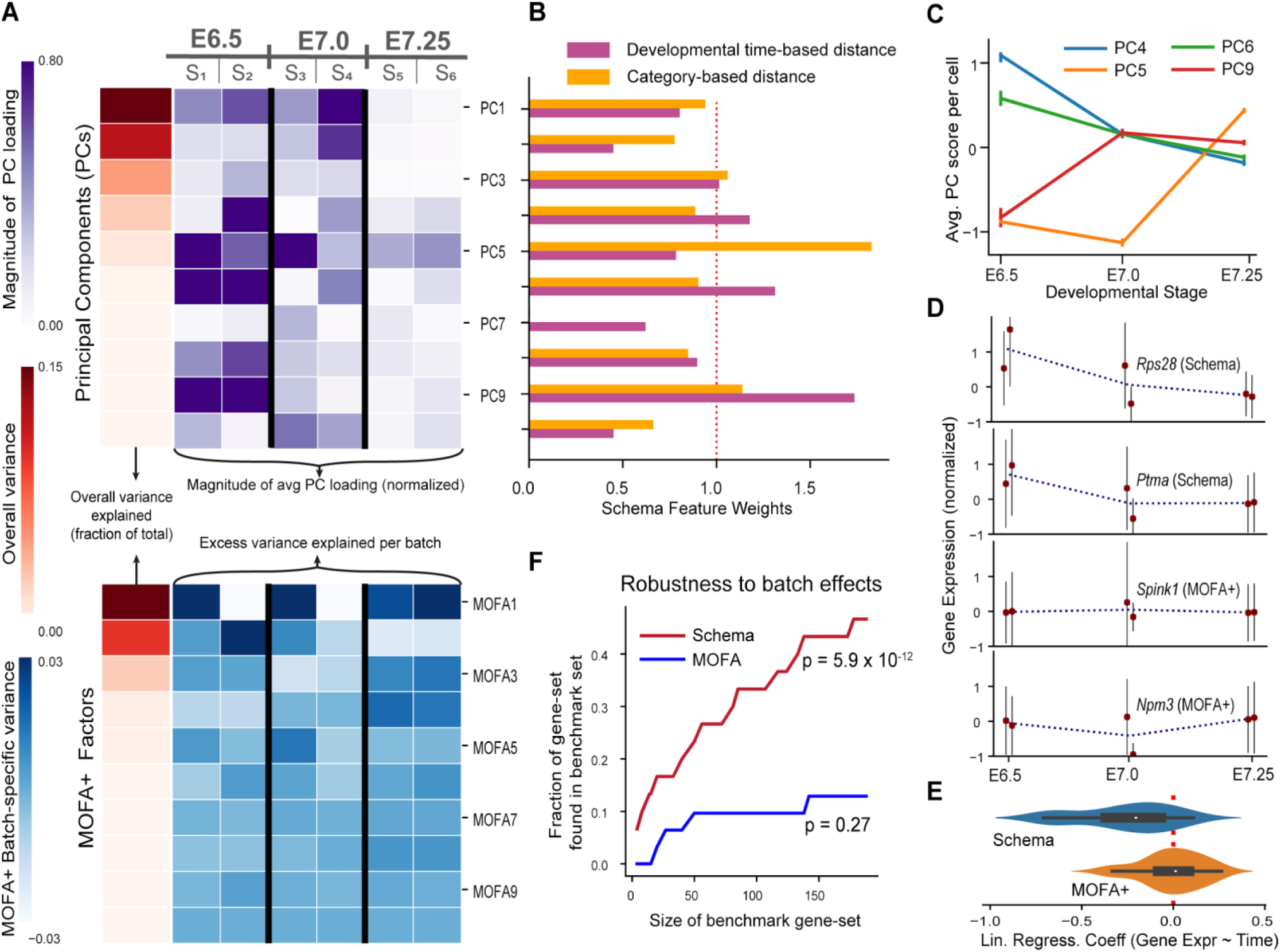
Batch-effect adjusted identification of differentially expressed genes along a developmental time course. (**A**) We obtained a dataset of developing mouse epiblast cells spanning three timepoints, with two experimental batches per timepoint. PCA and MOFA+ components show significant within-timepoint variability. In this panel, loadings of each principal component (PC) were normalized to zero mean and unit standard deviation. (**B, C**) Weights computed by Schema after accounting for batch effects and developmental age with two different distance metrics, one that provides Schema with temporal-ordering and another that does not provide this order. When incorporating order information, Schema down-weights PC5, which shows substantial within-timepoint, batch-related variability, and up-weights PC9, which has higher correlation with time. Correspondingly identified PCs reflect the effect of these metric. (**D, E**) Schema identifies genes with monotonically changing expression. For each gene identified by Schema or MOFA+, we regressed its expression (normalized to zero mean and unit standard deviation) against developmental time, encoding stages E6.25, E7.0 and E7.25 as timepoints 1, 2 and 3, respectively. Consistent with stage-dependent monotonicity in expression, the fitted slopes for Schema genes were significantly different from zero (two-sided *t*-test, *p* = 3.83 x 10^−6^); this was not true of MOFA+ (*p* = 0.77). (**F**) Schema has stronger overlap with batch-effect adjusted benchmark sets of differentially expressed genes (hypergeometric test with Bonferroni correction, *p* = 5.9 x 10^−12^ for the benchmark set of size 188).

We evaluated Schema alongside MOFA+, a recently introduced single-cell multimodal analysis technique^21,53^. Schema and MOFA+ approach the data synthesis problem from complementary perspectives: while the emphasis in Schema is to identify important features of the primary dataset and its corresponding transformation that reflects a synthesis of the various modalities, MOFA+ focuses on *de novo* identification of features that explain the covariation across modalities. In Argelaguet et al.’s MOFA+ analysis of this dataset, the authors identified 10 factors that capture similar information to the top principal components (**Supp Figure 5**). To identify differentially expressed genes with MOFA+, we selected the top genes from two factors (MOFA1 and MOFA4) reported by Argelaguet et al. as capturing developmental variation (**Methods**).

In addition to accounting for batch effects, we could also configure Schema to reduce the weight of transient changes in expression, thus identifying genes with monotonically changing expression along the time course (**Figure 3B-D**). To do so, we encoded developmental age as a distance metric by specifying zero distance between cells at the same timepoint, unit distance between directly adjacent timepoints, and an additive sum of the unit distances across more separated timepoints. As a control, we also tested a metric that did distinguish between the stages but did not increase in time, finding that the highest-weighted feature (PC5) in that case was indeed non-monotonic (**Methods**, **Figure 3B-C**). To encode batch effect as a distance metric, we specified zero distance between cells in the same replicate and unit distance otherwise.

We estimated the set of differentially expressed genes as the top-loading genes of the principal components up-weighted by Schema. Seeking to evaluate if the Schema or MOFA+ genes did show time-dependent monotonicity in expression, we linearly regressed each identified gene’s normalized expression against an ordering of the three developmental stages (**Methods**). We found that the Schema genes corresponded to regression coefficients significantly different from zero (**Figure 3D-E**), consistent with time-dependent monotonicity (two-sided *t*-test, *p* = 3.83 x 10^−6^); this was not true of MOFA+ (*p* = 0.77).

Next, we evaluated the batch-effect robustness of Schema and MOFA+ gene sets. Our configuration of Schema balances batch-effect considerations against differential expression considerations. For instance, introducing the batch-effect objective in Schema reduces the weights associated with the first and second principal components (PC1 and PC2), which show substantial within-timepoint batch-effect variations without a compensating time-dependent monotonicity, by 11% and 17%, respectively. In comparison, explicitly up-weighting “good” variation or down-weighting “bad” variation is difficult when using MOFA+. To systematically evaluate the batch-effect robustness of Schema and MOFA+ gene sets, we constructed benchmark sets of differentially expressed genes by applying a standard statistical test, adjusting for batch effects by exploiting the combinatorial structure of this dataset. Specifically, we aggregated over computations that each considered only one replicate per timepoint (**Methods**). We then measured the overlap of Schema and MOFA+ gene sets with these benchmarks (**Figure 3F**) and found that, compared to MOFA+, the Schema gene set shows a markedly higher overlap with the benchmarks that is statistically significant (hypergeometric test with Bonferroni correction, *p* = 5.9 x 10^−12^ for the benchmark set of size 188). Schema allows us to express the intuition that variation attributable to batch effects should be ignored while variation attributable to developmental age should be highlighted.

### Spatial density-informed differential expression among cerebellar granule cells

We performed some preliminary analysis with Schema of spatial transcriptomics data, another increasingly important multimodal scenario, here encompassing gene expression, cell-type labels, and spatial location. We obtained Slide-seq data containing 62,468 transcriptomes that are spatially located in the mouse cerebellum. In the original study, these transcriptomes were assigned to putative cell types (noting that these transcriptomes are not guaranteed to be single-cell), and thus cell types are located throughout the tissue ^10,26^. Interestingly, we observed spatial density variation for certain cell types; specifically, transcriptomes corresponding to granule cell types are observed in regions of both high and low spatial density (**Supp Figure 1B** in this paper; also Figure 2B of Rodriques et al. ^10^). We therefore reasoned that we could use Schema to identify genes that are differentially expressed in granule cells in high density areas versus granule cells in low density areas. Schema is well suited to the constrained optimization setting of this problem: we want to optimize for genes expressed specifically in granule cells *and* in dense regions, but not all granule cells are in dense regions and not all cells in dense regions are granule cells. We specified RNA-seq data as the primary modality and spatial location and cell-type labels as the secondary modalities, with spatial density controlled by a distance metric that scores two cells as similar if their spatial neighborhoods have matching densities (**Methods**). The densely-packed granule cell genes identified by Schema are strongly enriched for GO terms and REACTOME pathways^27^ related to signal transmission, e.g., ion-channel transport (REACTOME FDR q = 1.82 x 10^−3^), ion transport (GO:0022853, FDR q = 1.8 x 10^−17^), and electron transfer (GO:009055, FDR q = 2.87 x 10^−11^). This finding suggests potentially greater neurotransmission activity within these cells (**Supp Table 1–2**, **Supp Figure 3–4, Supp Text 4**).

### Schema outperforms alternative methods for spatial transcriptomic analysis

We sought to benchmark our method by comparing the robustness of Schema’s results with those based on canonical correlation analysis (CCA) and with two methods specifically intended for spatial transcriptomics, namely SpatialDE^18^ and Trendsceek ^19^.

An important point is that CCA, SpatialDE, and Trendsceek are less general than Schema and therefore require non-trivial modifications to approximately match Schema’s capabilities. CCA is limited in that it can correlate only two datasets at a time, whereas here we seek to synthesize *three* modalities: gene expression, cell-type labels, and spatial density. We adapted CCA by correlating two modalities at a time and combining the sub-results (**Methods**). In the case of SpatialDE and Trendsceek, their unsupervised formulation does not allow the researcher to specify the spatial features to pick out (we focus on spatial *density* variation). To adapt these, we collated their results from separate runs on granule and non-granule cells (**Methods**). Notably, the *ad hoc* modifications required to extend existing methods beyond two modalities underscore the benefit of Schema’s general analytic formulation that can be naturally extended to incorporate any number of additional data modalities.

To evaluate the stability and quality of spatial transcriptomic analysis across different techniques, we analyzed three replicate samples of mouse cerebellum tissue (coronal sections prepared on the same day^10^; pucks 180430_1, 180430_5, 180430_6) and compared the results returned separately for each replicate (**Methods**). While both Schema and CCA identify a gene set that ostensibly corresponds to granule cells in dense regions (**Supp Figure 1D** and **Supp Figure 3**), the gene rankings computed by Schema are more consistently preserved between pairs of replicates than those computed by CCA, with the median Spearman rank correlation between sample pairs being 0.68 (Schema) versus 0.46 (CCA). Likewise, with Schema, 69.1% of enriched GO biological-process terms are observed in all three samples and 78% are in at least two samples. The corresponding numbers for CCA were 35.7% and 59.5%, respectively (FDR *q* < 0.001 in all cases). We therefore find that Schema’s results are substantially more robust across the three replicates. We also find that Schema, in not seeking an unconstrained optimum, is more robust to overfitting to sample-specific noise than CCA (**Methods; Supp Figure 1E**).

When performing the same gene list robustness analysis with SpatialDE and Trendsceek, while also looking at the stability of their gene rankings specific to the precursor cell type (gray points in **Supp Figure 1E**), we found that SpatialDE produced slightly more stable gene rankings than Trendsceek, with median sample-pair correlations of 0.089 and −0.002, respectively, but these were still much lower than those for Schema. We also observed that SpatialDE and Trendsceek had substantially longer running times and we performed our analysis of the two methods on subsets of the overall dataset (see “Schema can scale to massive single-cell datasets” for precise runtime and memory usage). These results demonstrate the robustness and efficiency of Schema’s supervised approach.

### Genomic location and accessibility inform variability in expression

We next sought to demonstrate the flexibility of Schema to analyses beyond cell type clustering and differential expression analysis. We turned to a study that simultaneously profiled gene expression and chromatin accessibility from 3,260 human A549 cells ^6^. Using Schema, we characterized the genomic regions (relative to a gene’s locus) where chromatin accessibility strongly correlates with the gene’s expression variability, i.e., regions whose accessibility is differentially important for highly variable genes. Schema assigned the highest weight to features associated with chromatin accessibility over long ranges, i.e., ~10 megabase (Mb) (**Supp Figure 2C**). Searching for an explanation, we investigated if highly variable genes share genomic neighborhoods and mapped gene loci to topologically associated domains^32^ (TAD) of this cell type. We found strong statistical evidence that highly variable genes are indeed clustered together in TAD compartments (**Supp Figure 2D**), supporting our findings from Schema and suggesting an epigenetic role in controlling gene expression variability.

This demonstration illustrates how Schema’s generality facilitates innovative explorations of multimodal data beyond, for example, cell type clustering or differential gene expression analysis. Here we chose genes to be the unit of observation, allowing us to design a primary dataset that links each gene’s RNA-seq measurements to ATAC-seq peak counts in its neighborhood. Specifically, each primary feature corresponds to a genomic window relative to the gene’s locus and is scored as the sum over all cells of peak counts in the window, each cell weighted by the gene’s expression. We created multiple features, each corresponding to a specific window size and placement (**Methods**, **Supp Figure 2B**). Reusing RNA-seq as the secondary modality, we designed a distance metric that captures similarity in expression variability: for each gene, we normalize and sort the vector of its expression values across cells so that identical vectors imply an identical pattern of gene expression variation (**Methods**). We used Schema as a feature-selection mechanism, where up-weighted features correspond to genomic windows important for explaining gene expression variability.

We then benchmarked this feature selection approach against a ridge regression^54^ where features of the primary modality were specified as the explanatory variables and the standard deviation of each gene’s expression (summarized from the secondary modality) was the response variable. Both analyses agreed in assigning the highest weights to features corresponding to long-range (~10 Mb) genomic regions upstream and downstream of a gene. However, Schema’s regularization mechanism helps produce more consistent and stable feature weights, as evaluated on subsets of genes grouped by strand orientation or chromosome (**Supp Figure 2C** and **Supp Table 3**).

Our finding that chromatin accessibility at a long range (~10 Mb) may play a role in gene expression variability aligns with the observation by Kim et al.^33^ that long-range DNA methylation predicts gene expression. To further investigate the connection between chromatin accessibility and gene expression variability, we analyzed gene membership in TADs, hypothesizing that gene expression variability is likely to be influenced by the organization of TADs in the genome. We analyzed the clustering of highly variable genes (HVGs) on TADs within A549 cells (inferred from Hi-C data, accession ENCFF336WPU in ENCODE^34,35^) and found that HVGs are indeed more likely to be clustered together in TADs (**Supp Figure 2D**). By two independent permutation-based tests (**Methods**), we were able to reject the null hypothesis that genes in the top quartile of variability are dispersed randomly across TADs (*p* < 0.004 in both cases, with Bonferroni correction). Schema-based feature weighting therefore revealed an association between genomic architecture and gene variability. Notably, these results show how synthesis of multimodal RNA- and ATAC-seq data not only benefits standard analyses like cell type inference, but also enables creative and diverse exploratory data analysis.

### Schema reveals CDR3 segments crucial to T-cell receptor binding specificity

To further demonstrate the generality of Schema, we applied it to synthesize data modalities beyond gene expression. We integrated single-cell multimodal proteomic and functional data with Schema to better understand how sequence diversity in the hypervariable CDR3 segments of T-cell receptors (TCRs) relates to antigen binding specificities^37^. *De novo* design of TCRs for an antigen of interest remains a pressing biological and therapeutic goal ^40,41^, making it valuable to identify the key sequence locations and amino acids that govern the binding characteristics of a CDR3 segment. Towards this end, we analyzed a single-cell dataset that recorded clonotype data for 62,858 T-cells and their binding specificities against a panel of 44 ligands^5^, and used Schema’s feature-selection capabilities to estimate the sequence locations and residues in the CDR3 segments of α and β chains important to binding specificity.

To estimate location-specific selection pressure, we ran Schema with the CDR3 peptide sequence data as the primary modality and the binding specificity information as the secondary modality, performing separate runs for α and β chains. In the primary modality, each feature corresponds to a CDR3 sequence location and we used the Hamming distance metric between observations (i.e. the number of locations at which two sequences differ, see **Methods**). Schema assigned relatively low feature weights to the location segments 3-9 (in α chain CDR3) and 5-12 (in β chain CDR3), suggesting those regions can tolerate greater sequence variability while preserving binding specificity.

To evaluate these results, we compared them to estimates based on CDR3 sequence motifs sourced from VDJdb^38^, a curated database of TCRs with known antigen specificities. In VDJdb, TCR motifs are scored using an adaptation of the relative-entropy algorithm^39^ by Murugan et al. that assigns a score for each location and amino acid in the motif. We aggregated these scores into a per-location score (**Methods**), allowing a comparison with Schema’s feature weights (**Figure 4**). While the comparison at locations 11-20 is somewhat complicated by VDJdb having fewer long sequences (**Methods**), there is agreement between Schema and VDJdb estimates on locations 1-10 where both datasets have good coverage (Spearman rank correlations of 0.38 and 0.92 for the α and β chains, respectively; **Figure 4C-D**). We note that weight estimation using Schema required only a single multimodal dataset; in contrast, extensive data collection, curation, and algorithmic efforts underlie the VDJdb annotations. The latter covers multiple experimental datasets, including the 10x Genomics dataset^5^ we investigated here; we saw similar results when comparing against an older version of VDJdb without this dataset.

**Figure 4:**
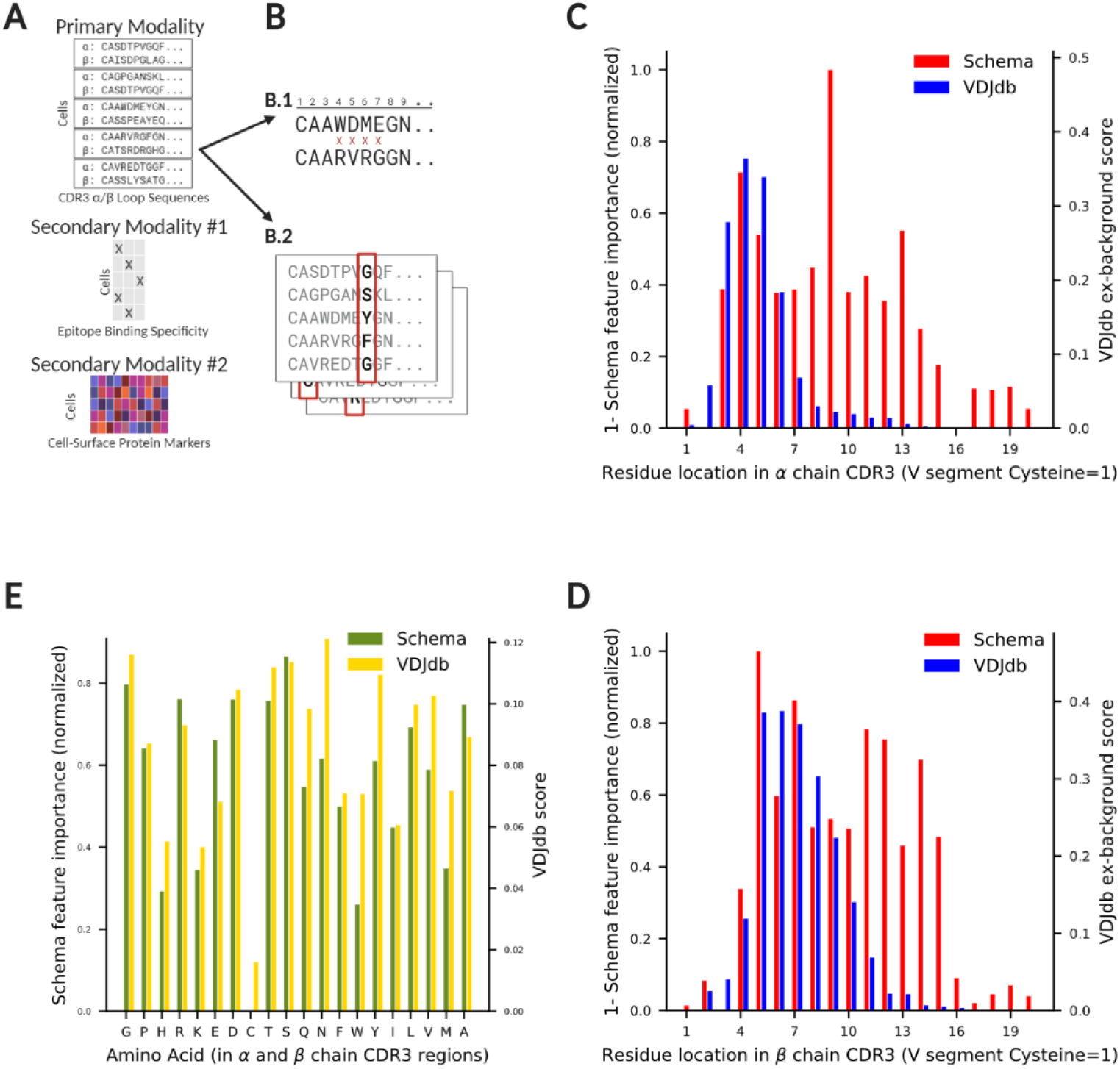
Schema reveals the locations and amino acids important in preserving binding specificity of T-Cell receptor CDR3 regions. (**A**) We analyzed a multi-modal dataset from 10x Genomics^5^ to understand how a T-cell receptor’s binding specificity relates to the sequence variability in the CDR3 regions of its α and β chains. The primary modality consisted of CDR3 peptide sequence data which we correlated with the secondary modality, the binding specificity of the cell against a panel of 44 epitopes. We optionally synthesized an additional modality, proteomic measurements of 12 cell-surface marker proteins, as a use-case of incorporating additional information (**Methods**). (**B**) We performed two Schema analyses: (*B.1*) To infer location-wise selection pressure, the feature vector of the primary modality was the CDR3 sequence, with the Hamming distance between two sequences as the metric. (*B.2*) The second analysis aimed to understand amino acid selection pressure at locations that showed high variability. For each such location, a one-hot encoding of the amino acid at the location was used as the feature vector; we operated an ensemble of Schema runs across various locations. (**C, D**) Schema identifies sequence locations 3-9 (α chain) and 5-12 (β chain) as regions where sequences can vary with a comparatively modest impact on binding specificity. We compared Schema’s scores to statistics computed from motifs in VDJdb. Here, we have inverted the orientation of Schema’s weights to align them with the direction of VDJdb weights. (**E**) Schema and VDJdb agree on the relative importance of amino acids in preserving binding specificity (Spearman rank correlation = 0.74, two-sided *t*-test *p* = 2 x 10^−4^). The low weight assigned to Cysteine is likely due to its infrequent occurrence, making the estimate unreliable.

Next, we used Schema to investigate the selection pressure on amino acids present in the variability-prone locations identified above (**Methods**). We first selected a sequence location (e.g., location 4 in α chain CDR3) and constructed a primary modality where each cell was represented by a one-hot encoding of the amino acid at the location (i.e., a 20-dimensional boolean vector). The secondary modality was binding specificity information, as before. We performed separate Schema runs for each such location of interest on the two chains, computing the final score for each amino acid as the average score across these runs. These scores are in good agreement with the corresponding amino acid scores aggregated from the VDJdb database (Spearman rank correlation = 0.74, two-sided *t*-test *p* = 2 x 10^−4^). The residue and location preferences estimated here can directly be used in any algorithm for computational design of epitope-specific CDR3 sequences to bias its search towards more functionally plausible candidate sequences.

Schema’s ability to efficiently synthesize arbitrarily many modalities, with their relative importance at the researcher’s discretion, allows information that might otherwise be set aside (e.g., metadata like batch information, cell line, or donor information) to be effectively incorporated, enhancing the robustness and accuracy of an analysis. In **Methods**, we exemplify this use-case on the TCR dataset by incorporating measurements of cell-surface markers as an additional secondary modality, hypothesizing that cell-surface protein levels should be unrelated to V(D)J recombination variability.

### Schema highlights secondary patterns while preserving primary structure

Another powerful use of Schema is to infuse information from other modalities into RNA-seq data while limiting the data’s distortion so that it remains amenable to a range of standard RNA-seq analyses. Having demonstrated this capability for cell type inference, we now explore another use case. Since widely-used visualization methods such as UMAP^42^ do not allow a researcher to specify aspects of the underlying data that they wish to highlight in the visualization, we sought to apply Schema to improve the informativity of single-cell visualizations. We leveraged Schema to highlight the age-related structure in an RNA-seq dataset of *Drosophila melanogaster* neurons^3^ profiled across a full lifespan, while still preserving most of the original transcriptomic structure. We chose RNA-seq as the primary modality and temporal metadata as the secondary modality, configuring Schema to maximize the correlation between distances in the two while constraining the distortions induced by the transformation (**Methods**). We then visualized the transformed result in two dimensions with UMAP.

While some age-related structure does exist in the original data, Schema-based transformation of the data more clearly displays a cellular trajectory consistent with biological age (**Figure 5**). Importantly, accessing this structure required only a limited distortion of the data, corresponding to relatively high values (≥ 0.99) of the minimum correlation constraint (**Figure 5C**). To verify that Schema was able to infuse additional age-related structure into RNA-seq data, we performed a diffusion pseudotime analysis of the original and transformed datasets and found that the Spearman rank correlation between this pseudotime estimate and the ground-truth cell age increased from 0.365 in the original data to 0.405 and 0.436 in the transformations corresponding to minimum correlation constraints of 0.999 and 0.99, respectively. In contrast, an unconstrained synthesis by CCA leads to a lower correlation (0.059) than seen in the original RNA-seq dataset; the corresponding CCA-based UMAP visualization is also less clear in conveying the cellular trajectory (**Supp Figure 10**). Schema thus enables visualizations that synthesize biological metadata, while preserving much of the distance-related correlation structure of the original primary dataset. With Schema, researchers can therefore investigate single-cell datasets that exhibit strong latent structure (e.g., due to secondary metadata like age or spatial location), needing only a small transformation to make that structure visible.

**Figure 5:**
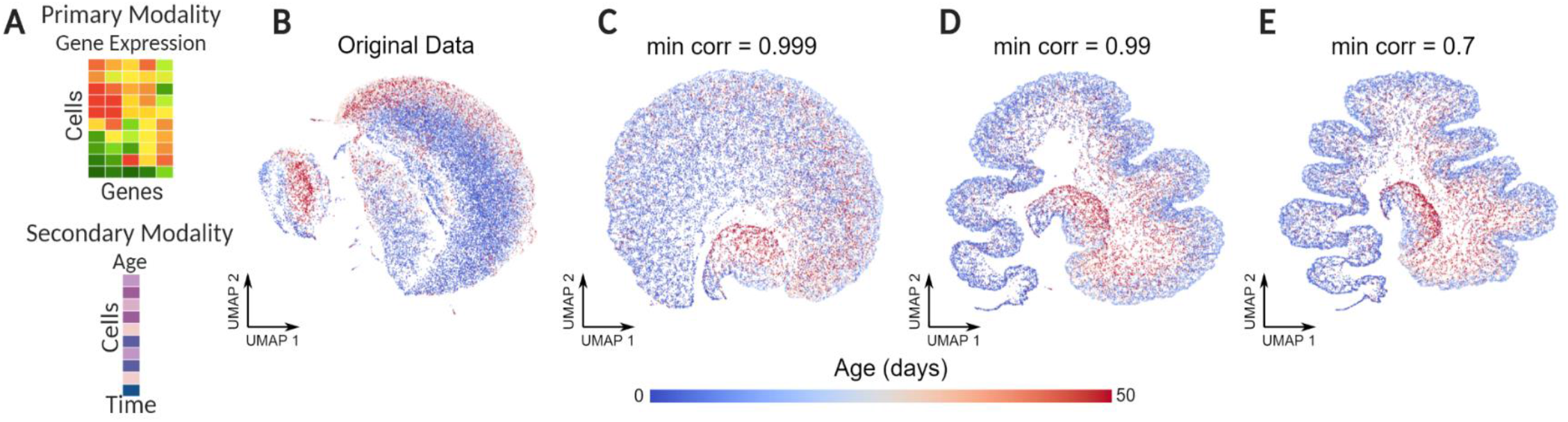
Incorporating temporal metadata into UMAP visualizations of aging neurons captures developmental changes. UMAP visualization of RNA-seq profiles of *D. melanogaster* neurons at 0, 1, 3, 6, 9, 15, 30, and 50 days after birth, representing the full range of a typical *D. melanogaster* lifespan. (**A**) The transcriptomic data (primary modality) was transformed to a limited extent using Schema by correlating it with the temporal metadata (secondary modality) associated with each cell. (**B**) UMAP visualization of the original transcriptomic data. (**C, D, E**) Visualizations of transformed data with varying levels of distortion. As the value of the minimum correlation constraint *s* approaches 1, the distortion of the original data is progressively limited. Decreasing *s* results in a UMAP structure that increasingly reflects an age-related trajectory.

### Schema can synthesize spliced and unspliced RNA counts to accentuate cell differentiation

We next leveraged the flexibility of Schema to study cell differentiation by synthesizing spliced and unspliced mRNA counts in a dataset of 2,930 mouse dentate gyrus cells^57^. Specifying spliced counts as the primary dataset and unspliced counts as the secondary dataset, we configured Schema to compute a transformation of the spliced data that maximizes the correlation of its Euclidean distances with those in the unspliced dataset while distorting the former only minimally (**Figure 6A-C**).

**Figure 6:**
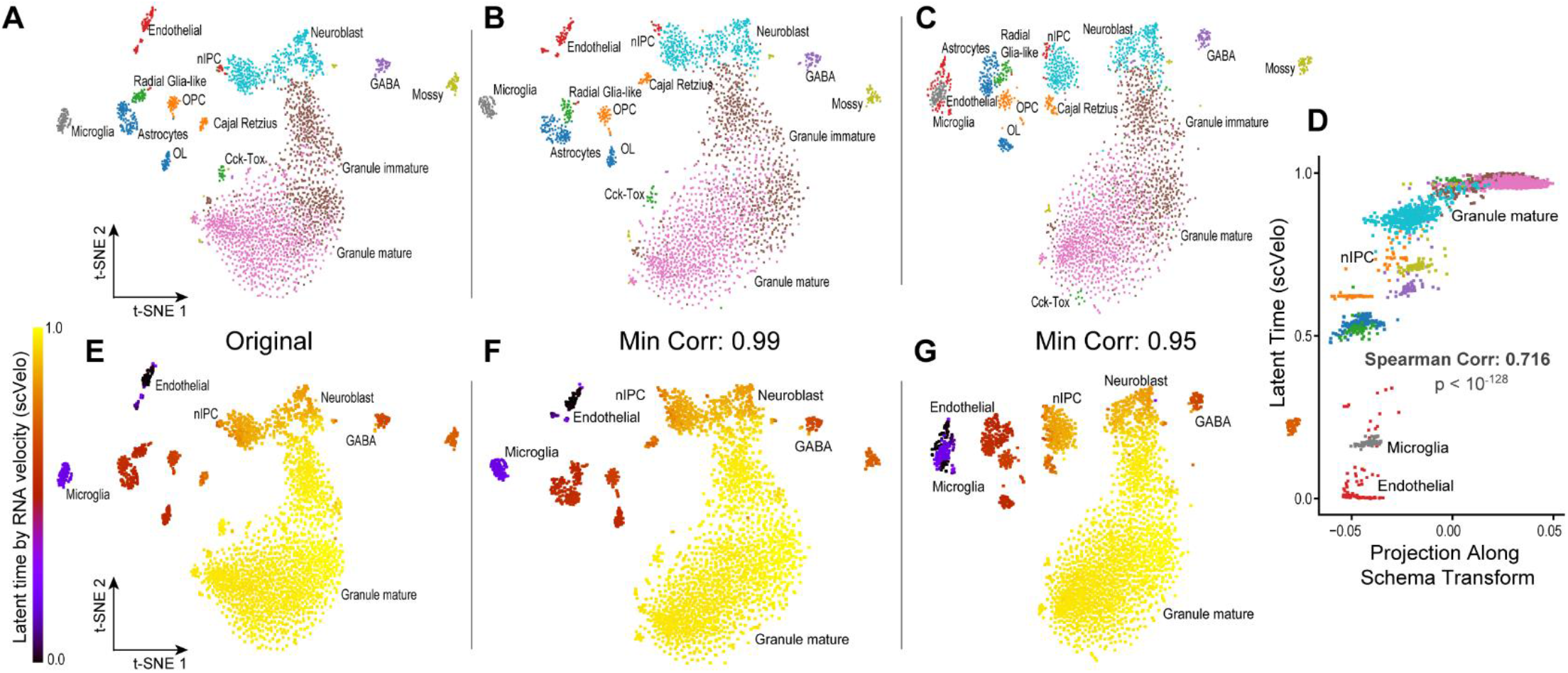
Synthesis of spliced and unspliced mRNA counts recovers RNA velocity and enables informative visualization. (**A**): t-SNE visualization of the spliced mRNA counts (**B-C**): We synthesized spliced and unspliced mRNA counts, with the former as the primary and the latter as the secondary modality, respectively. Schema’s transformation picks up the time derivative of gene expression, thus accentuating the cell differentiation process. t-SNE visualizations of synthesized data with 0.99 and 0.95 minimum correlation, respectively, are shown. (**D**) Schema’s results are in agreement with the RNA velocity tool, scVelo. By measuring each cell’s Schema transformation, we computed a pseudotime estimate which we found to be significantly correlated with scVelo’s latent-time estimate (Spearman rank correlation = 0.716, two-sided t-test *p* < 10^−128^). (**E-G**): Same t-SNE visualizations as above, but with cells colored by their scVelo latent-time, showing that Schema puts cells at similar differentiation stages progressively closer.

Our intuition here is the same as that underlying RNA velocity techniques: correlating spliced and unspliced counts in a cell should pick up on the time derivative of a cell’s expression state and thus illuminate the cell differentiation process. To validate this intuition, we computed a pseudotime measure from the difference between transformed and original RNA-seq data, finding it to be highly correlated with the latent-time estimate produced by Bergen et al.’s RNA velocity tool scVelo^6^ (Spearman rank correlation 0.72, two-sided *t*-test *p* < 10^−128^, **Figure 6D**, **Methods**). Since Schema relies on the same underlying biological phenomena as specialized RNA velocity tools but analyzes the data differently, these results show the breadth of Schema’s generality and may be used to help supplement and strengthen the findings from standard RNA velocity analysis.

Schema can complement methods like scVelo by facilitating additional analyses. As in our demonstrations of cell type inference and UMAP visualization, the transformed data produced here by Schema incorporates additional information (the time derivative of expression) but remains analyzable as an RNA-seq dataset. As an example, we visualized the transformed dataset with t-SNE, finding that the two-dimensional t-SNE plot of the Schema-transformed data places more closely together cell types at similar stages of differentiation (as quantified by scVelo latent-time, **Figure 6E-G**). To confirm this visual observation, we computed the Spearman rank correlation of scVelo latent-time differences between pairs of cells and their corresponding Euclidean distances in the t-SNE embedding space, finding that it increases from 0.397 in the original dataset to 0.432 in the transformation corresponding to a minimum correlation constraint of 0.95 (**Methods**). In contrast, an unconstrained synthesis using CCA produced a substantially lower correlation of 0.163; see **Supp Figure 10** for the corresponding CCA-based t-SNE visualization. Schema can thus facilitate visualizations that reflect the deeper underlying differentiation processes.

### Schema can scale to massive single-cell datasets

We have designed Schema to process large single-cell datasets efficiently, with modest memory requirements. On average, Schema processes data from a Slide-seq replicate (three modalities, 20,823 transcriptomes x 17,607 genes) in 34 minutes, requiring less than 5GB of RAM in the process (**Supp Table 4**). The runtime includes the entire set of Schema sub-runs performed over an ensemble of parameters, as well as the time taken for the pre-processing transformation.

Schema’s efficiency stems from our novel mathematical formulation. Deviating from standard metric learning approaches, we formulate the synthesis problem as a quadratic-program optimization, which can be solved much faster than the semi-definite program formulations typically seen in these approaches (**Supp Text 1**). Additionally, while the full Schema algorithm has quadratic scalability in the number of cells, our formulation allows us to obtain good approximations with *provably* bounded error using only a logarithmic subsample of the dataset (**Supp Text 3**), enabling *sublinear* scalability in the number of cells that will be crucial as multimodal datasets increase in size.

## Discussion

We designed Schema to be a powerful approach to multimodal data analysis. Schema is based on an elegant conceptual formulation in which each modality is defined using a distance metric. The strength of this intuition enables analysis of an arbitrary number of modalities and applicability to any modality, so long as it is possible to define an appropriate distance metric.

Our approach is supervised, allowing the researcher to control which modality to transform, the amount of distortion, and the desired level of agreement between modalities. While existing methods like Seurat v3^16^ and LIGER^17^ are designed for unsupervised discovery of common patterns across experiments, Schema’s supervised formulation facilitates a broader set of investigations, enabling us to not only infer cell types and identify gene sets but also rank amino acids by selection pressure and estimate the relative importance of different genomic neighborhoods in epigenetic regulation.

When choosing a primary (i.e. reference) modality, we recommend selecting the most high-confidence modality or the one for which feature selection will be most informative. In many of our demonstrations, we chose RNA-seq as the primary modality since it is often the modality where preprocessing and normalization are best understood, boosting our confidence in it; additionally, transformed RNA-seq data lends itself to a variety of downstream analyses. Once a primary modality has been designated, Schema can synthesize an arbitrary number of secondary modalities with it. In contrast, methods designed around pairwise modality comparison need *ad hoc* adaptations to accommodate additional modalities. Schema’s approach is advantageous not only for datasets with more than two modalities^5,44^ but also in cases where metadata (e.g. batch information and cell age) can be productively incorporated as additional modalities.

Intuitively, our correlation-based alignment approach has parallels to kernel canonical correlation analysis (kernel CCA), a generalization of CCA where arbitrary distance metrics can be specified when correlating two datasets. While Schema offers similar flexibility for secondary modalities, it limits the primary modality to Euclidean distances. Introducing this restriction enhances scalability, interpretability and robustness. Unlike kernel CCA, the optimization in Schema operates on matrices whose size is independent of the dataset’s size, enabling it to scale sub-linearly to massive single-cell datasets. Also, the optimal solution is a scaling transform that can be naturally interpreted as a feature-weight vector. Perhaps most importantly, Schema differs from kernel CCA in performing a constrained optimization, thus reducing the distortion of the primary dataset and ensuring that sparse and low-confidence secondary datasets do not drown out the primary signal.

The constrained optimization in Schema acts as regularization, helping ensure that the computed transformation and feature selection remain biologically meaningful. By choosing a high-confidence modality as the primary modality and bounding its distortion when incorporating the secondary modalities, Schema enables information synthesis while retaining high-confidence insights. This bound on the distortion is an important parameter, directly controlling how much the secondary modalities inform the primary dataset; values approaching 1 will increasingly limit the influence of the secondary modalities. Therefore, we recommend that studies using Schema for feature selection should aggregate the results over a range of values of this parameter while analyses that utilize only a single parameter should keep it high (≥ 0.9, the default setting in our implementation is 0.99) to preserve fidelity with the original dataset. We also strongly recommend that studies with a single parameter should report the value of this parameter alongside their results.

Interesting future methodological work could explore alternative formulations of the Schema objective, potentially including more complex nonlinearities than our quadratic-program formulation. Schema can also guide further biological experiments that profile only the highly-weighted features based on other data modalities, enabling efficient, targeted follow-up analysis.

Given the current pace of biotechnological development, we anticipate that high-throughput experiments, and their conclusions, will increasingly rely on more than one data modality, underscoring the importance of Schema and its conceptual framework. Schema is publicly available for use at http://schema.csail.mit.edu and as the Python package *schema_learn*.

## Methods

### Correlation-based alignment and quadratic programming optimization

Underlying our definition of the alignment of metrics is the intuitive notion that metrics are similar if the ordering of pairwise distances between the two metrics are close. A proxy for measuring this alignment is the Pearson correlation coefficient. For Schema, the goal is thus that pairwise distances in the transformed space are highly correlated with pairwise distances under each metric.

One of the advantages of the Pearson correlation coefficient is that it is amenable to optimization via *quadratic programming* (QP). QP is a generalization of linear programming, allowing a quadratic objective function. We learn a *scaling transformation u* (**Supp Text 1**) on the primary dataset *X* such that the pairwise distances of the transformation *u* * *x_i_* (where * denotes coordinate-wise multiplication, for each *x_i_* ∈ *X*) are highly correlated with the pairwise distances in the secondary modalities. We codify our intuition of the importance of the primary dataset by ensuring that the correlation of transformed pairwise distances with the original dataset is higher than some researcher-specified threshold. The scaling transformation has the appealing property of being interpretable as a *feature selection*. The higher the coordinate *u_i_*, the more important that coordinate is for alignment. Thus, by selecting the top coordinates by their weights, we can access the genes most important for aligning the modalities.

### Mathematical formulation

Suppose we have *N* observations across *r* datasets *D_j_*, *j* = 1,2, …, *r*, where 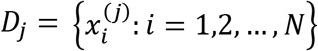 contains data (categorical or continuous) for each observation. We will refer to *D*_1_ as the *primary* dataset and the rest as secondary. Each dataset’s dimensionality and domain may vary. In particular, we assume *D*_1_ is *k*-dimensional. Each dataset *D_j_* should also have some notion of distance between observations attached to it, which we will denote *ρ_j_*, so 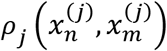 is the distance between observations *n* and *m* in *D_j_*. Since our entire framework below deals in *squared* distances, for notational convenience we will let *ρ_j_* be the squared distances between points in *D_j_*; also, we drop the superscript in 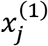 when referring to the primary dataset *D*_1_ and its data.

The goal is to find a transformation *Ω* with *Ω*(*D*) generating a dataset *D*\* such that the Euclidean metric *ρ*\* on *D*\* aligns the various metrics *ρ_j_*, each informed by its respective modality. Non-Euclidean primary distance metrics *ρ*\* are allowed if they can be computed as a sum of *k* terms, one for each feature (e.g. Hamming distance). We emphasize that none of the secondary *ρ_j_* need to be Euclidean. This setup is quite general, and we now specify the form of the transformation *Ω* and the criteria for balancing information from the various metrics. Here, we limit *Ω* to a *scaling transform*. That is, *Ω*(*D*) = {*diag*(*u*)*x*: *x* ∈ *D*} for some *u* ∈ *R^k^* and *diag*(*u*) is a *k* × *k* diagonal matrix with *u* as its diagonal entries. Then, the squared distance between points under the transformation is given by:

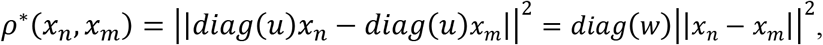

where *w* is the element-wise square of *u*, i.e. 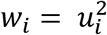. The scaling transform *u* acts as a feature-weighting mechanism: it chooses the features of *D*_1_ that align the datasets best (i.e. *u_i_* being large means that the *i*th coordinate of *D*_1_ is important). We note here that a natural extension would be allowing *general linear* transformations for *Ω*; however, in that context, the fast framework of quadratic programming would need to be substituted for the much slower framework of semidefinite programming.

Here, our approach to integration between the metrics *ρ_j_*. is to learn a metric *ρ*\* that aligns well with all of them. Our measure of the alignment between *ρ*\* and *ρ_j_* is given by the Pearson correlation between pairwise squared distances under two metrics. Intuitively, maximizing the correlation coefficient encourages distances under *ρ*\* to be large when the corresponding *ρ_j_*. distances are large and vice versa. This can be seen from the expression

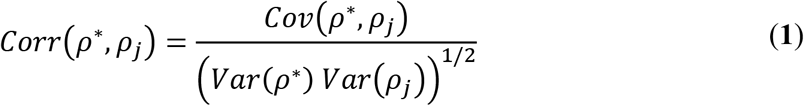

To deal with multiple modalities, we try to maximize the correlation between *ρ*\* and the distances on each of the metrics, allowing the user to specify how much each modality should be weighted. We also allow a hard constraint, whereby the correlation between the pairwise distances in the transformed data and in the primary dataset is lower-bounded. Our goal is thus to find

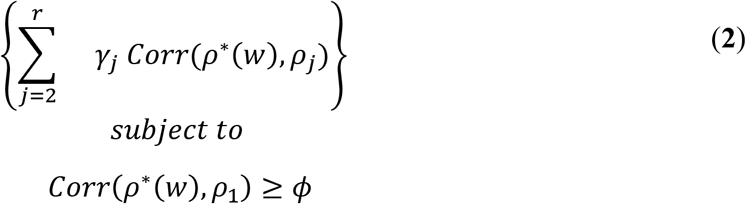

where *γ_j_* and *ϕ* are hyperparameters that determine the importance of the various metrics. We have also highlighted that *ρ*\* is a function of w and is determined entirely by the solution to (**2**). In the rest of our discussion, we will abuse notation and primarily use *w*, rather than *ρ*\*, to refer to the optimal metric. The machinery of *quadratic programming* makes this optimization feasible.

### Setting up the quadratic program

As motivated above, quadratic programming (QP) is a framework for constrained convex optimization problems that allows a quadratic term in the objective function and linear constraints. The general form is

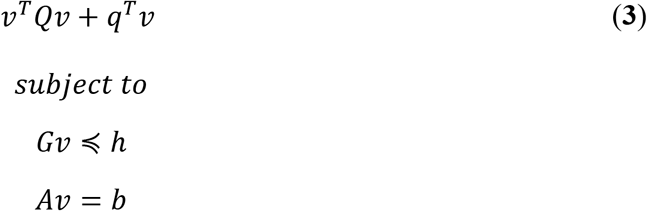

for given *Q, q, G, h, A*, and *b*, where *Q* is a positive semidefinite (psd) matrix and the notation ≼ means the inequality is true for each coordinate (i.e., *y* ≼ *z* means *y_i_* ≤ *z_i_* for all *i*).

To put our optimization (**2**) in a QP formulation, we expand the covariance and variance terms in the definition of correlation in (**1**), and show that the covariance is *linear* in the transformation and variance is *quadratic*:

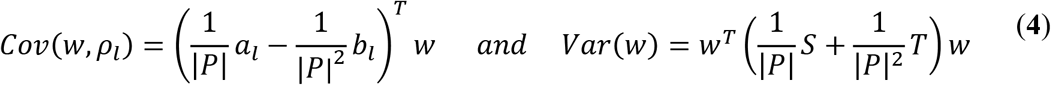

where *a_l_* and *b_l_* are *k*-dimensional vectors that depend only on *D_l_*; and *S* and *T* are *N* × *k* matrices that depend only on *D*_1_, and *P* is the set of pairs of observations, where |·| denotes set cardinality. It is also not hard to show that 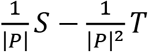 is psd, as required. For details of the derivation, see the **Supp Text 2**.

There is one more difficulty to address. The correlation is the *quotient* of the covariance and the standard deviation, and the QP framework cannot handle quotients or square roots. However, maximizing a quotient can be reframed as maximizing the numerator (the covariance), minimizing the denominator (the variance), or both.

We now have the ingredients for the QP and can frame the optimization problem as

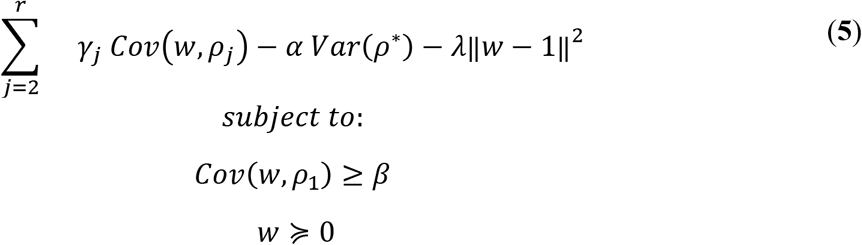

where 0 and 1 are the all-zeros and all-ones vectors (of the appropriate length) respectively. Here, *λ* is the hyperparameter for regularization of *w*, which we want to penalize for being too far away from the all-ones vector (i.e., equal weighting of all the features). One could also regularize the *l*_2_ norm of *w* alone (i.e., incorporate the term −*λ*∥*w*∥^2^), which would encourage *w* to be small; we have found that empirically the choices yield similar results.

This program can be solved by standard QP solvers (see **Supp Text 2**) for the full details of how to put the above program in canonical form for a solver), and the solution *w*\* can be used to transform unseen input data, using *u*\* ∈ *R^k^*, where 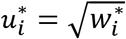.

### Hyperparameters

A well-known challenge for machine learning algorithms is interpretability of hyperparameters. Here, the QP solver needs values for *λ*, *α*, and *β*, and specifying these in a principled way is a challenge for users. Our approach is thus to allow the user to specify more natural parameters. Specifically, we allow the user to specify minimum correlations between the pairwise distances in *D*\* and the primary dataset *D*_1_. Formally, the user can specify *s* such that

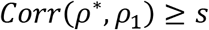

and *q* such that

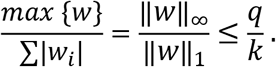

The quantity *q* thus controls the maximum weight that any one feature can take.

While these quantities are not directly optimizable in our QP formulation (**5**), we can access them by varying the hyperparameters *λ, α*, and *β*.

Intuitively, we note that the choice of *λ* controls whether *w* satisfies *q* and that *α* and *β* control whether the correlation constraint *s* is satisfied. To satisfy these constraints, we simply grid search across feasible values of {*λ*, *α*, *β*}: we solve the QP for fixed values of *λ*, *α*, and *β*, keeping only the solutions for which the {*s*, *q*} constraints are satisfied. Of these, we choose the most optimal. The efficiency of quadratic programming means that such a grid search is feasible, which gives users the benefit of more easily interpretable and natural hyperparameters.

### Recommendations for setting s and q

We recommend that only *s* (minimum correlation) and not *q* (maximum feature weight) be used to control Schema’s optimization. The default value of *q* in our implementation is set to be very high (10^3^) so that it is not a binding constraint in most cases. We recommend not changing it and in future versions of Schema we may reformulate the QP so that *q* is entirely removed. To limit the distortions in the primary modality, we recommend that *s* be set close to 1: the default setting of *s* is 0.99 and we recommended values ≥ 0.9. When Schema is used for feature selection, we recommend aggregating results across an ensemble of runs over a range of *s* values (a wide range is recommended here) to increase the robustness of the results.

### Preprocessing transforms

Standard linear decompositions, like PCA or NMF are useful as preprocessing steps for Schema. PCA is a good choice in this regard because it decomposes along directions of high variance; NMF is slower, but has the advantage that it is designed for data that is non-negative (e.g., transcript counts). Since the transform *ω* that we generate can be interpreted as a feature-weighting mechanism, we can identify the directions (in PCA) or factors (in NMF) most relevant to aligning the datasets. Here the user can employ arbitrary feature-sets including, for instance, a union of features from two standard methods (e.g., set-union of PCA and CCA features) or those generated by another single-cell analysis method like MOFA+^21^.

### Motivating the choice of correlation as an objective

As a measure of the alignment between our transformation and a dataset, correlation of pairwise distances is a flexible and robust measure. An expanded version of arguments in this paragraph is available in the appendices. Given a pair of datasets, the connection between their pairwise-distance Spearman rank correlation and the neighborhood-structure similarity is deep: if the correlation is greater than 1 – *ϵ*, the fraction of misaligned neighborhood-relationships will be less than 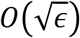. There is a *manifold* interpretation that is also compelling: assuming the high-dimensional data lies on a low-dimensional manifold, small Euclidean distances are more accurate than large distances, so the *local* neighborhood structure is worth preserving. We can show intuitively that optimizing the correlation aims to preserve local neighborhood structure. Using correlation in the objective also affords the flexibility to broaden *Corr*(*w*, *ρ_j_*) in (**2**) to any function *f_j_* of the metric, i.e., *Corr*(*w*, *f_j_* ∘ *ρ_j_*); this allows us to invert the direction of alignment or more heavily weigh local distances. As RNA-seq dataset sizes reach millions of cells, even calculating the *O*(*N*^2^) pairwise distances becomes infeasible. In this case, we sample a subset of the pairwise distances. As an estimator, sample correlation is a robust measure, allowing Schema to perform well even with relatively small subsets; in fact, we only need a sample size *logarithmic* in our desired confidence level to generate high-confidence results (**Supp Text 3**). This enables Schema to continue scaling to more massive RNA-seq datasets.

### Inference of cell types by synthesizing gene expression and chromatin accessibility

Applying the *TruncatedSVD* function in the Python library *scikit-learn*^54^ (version 0.23.1), we reduced the dimensionality of the primary (RNA-seq) and secondary (ATAC-seq) datasets to their top 100 and 50 components, respectively, and specified these as the inputs to Schema. We chose to perform SVD instead of PCA since only the former can work with sparse matrices (in particular, the ATAC-seq matrix had 11,296 rows and 247,293 columns). The minimum correlation threshold in Schema was set to 0.99 and Leiden clustering was performed with the Python package *leidenalg*^56^ (version 0.8.1) with partitioning of the neighbor graph based on the modularity measure.

We performed canonical correlation analysis (CCA) on the same dimensionality-reduced primary and secondary datasets as supplied to Schema and computed 30 CCA factors, performing Leiden clustering using these. To implement the heuristic approach described by Cao et al.^6^, we grouped the 11,296 cells into *k*=300 clusters by k-means clustering of RNA-seq data; results were robust to the choice of *k*. Each cluster (“pseudocell”) was represented by the average ATAC-seq profile of its member cells, with these aggregated profiles forming the input to the Leiden clustering algorithm.

### Differential expression analysis while accounting for batch effects

This mouse gastrulation dataset was originally described by Pijuan-Sala et al.^57^ and investigated by Argelaguet et al.^21,53^ using the MOFA+ algorithm. We operated on the data as preprocessed and made available by them and, for the MOFA+ evaluation in this paper, also used their pretrained models.

We first reduced the RNA-seq data (primary modality) to its top 10 principal components (PCs), to be in line with the 10 MOFA+ factors from Argelaguet et al. The MOFA+ algorithm can be thought of as a generalization of PCA and we did indeed observe that the top PCs were very similar to the top MOFA+ components (**Supp Figure 4**), validating that Schema was able to access the same sources of variation as found by MOFA+ here.

We configured Schema to use batch information as a secondary modality with weight −1 and developmental age information as a secondary modality with weight +1; thus, correlation with the former was minimized and the latter was maximized. The minimum correlation threshold was set to 0.9; we found that the results were robust to variations in this setting (0.8 and 0.95).

Since Schema accepts arbitrary distance measures on secondary datasets, we could investigate the impact of treating developmental timepoints as categories rather than a time ordering. In the categorical distance metric, we defined two cells to be at distance 0 if they were at the same developmental timepoint and at distance 1 otherwise. In the time-ordering metric, we specified the first and third time-points to be at distance 2 apart and the middle time-point to be at distance 1 from either end. The two distance measures lead to different feature-selection results from Schema, reflecting the distinct underlying variations in expression profiles. For category-based distances, PC5 receives the highest weight while PCs 4 and 6 are given higher weights in the time-ordering case (**Figure 3B**). This happens because the mean expression level of PC5 shows large variation across the three time-points but does not change monotonically along the time course; in contrast, the PCs 4 and 6 display expression profiles that change monotonically with developmental age (**Figure 3C**). Schema’s flexibility to incorporate a distance measure that highlights specific variability patterns can thus enable researchers to identify precisely targeted gene-sets.

To create a gene set from Schema’s feature weighting, we selected the intersection of *k* top loadings (by absolute value) of PCs up-weighted by Schema (PCs 4, 6 and 9 for the time-ordering metric); we choose *k* so that the intersection contained 30 genes. For MOFA+, we chose genes that had the top loadings (by absolute value) in the factors MOFA1 or MOFA4. Here, we were following Argelaguet et al who, after an investigation of the various MOFA+ factors, had identified these two as the most relevant to understanding developmental age variability. Since the top loadings of the two MOFA+ factors do not overlap much, we chose the top 16 genes from each, with their union consisting of 31 genes (there was one overlap between the two subsets).

For each gene identified by Schema or MOFA+, we regressed its expression against developmental time, encoding stages E6.25, E7.0 and E7.25 as timepoints 1, 2 and 3, respectively. The gene’s expression profile (across all cells) was first normalized to zero mean and unit standard deviation.

We created batch-effect adjusted benchmark gene sets by using different combinations of replicates. One can create a subset of the original dataset by sampling cells from only one of the two replicates at each time-point. By iterating over all possible combinations of replicates, we created eight such subsets. These subsets differ in the batch information they contain but share the same developmental age information. Using the Wilcoxon rank sum test in *scanpy*^36^, we identified genes differentially expressed between the first (E6.5) and last (E7.25) stage in each subset and defined the benchmark gene set to consist of genes that are differentially expressed across a majority of the subsets. The benchmark set is thus robust to batch effects, being comprised of genes whose differential expression stands out across different replicates (i.e. batches). By varying the thresholds of the test, we could create benchmark sets of varying sizes and measured the overlap of Schema and MOFA+ gene sets with these. The Schema gene set has a higher overlap and for benchmark sets of all sizes, its overlap with them was significant (hypergeometric test with Bonferroni correction, *p* = 5.9 x 10^−12^ for the benchmark set of size 188 and Bonferroni-corrected *p* < 10^−5^ for benchmark sets of all sizes, **Figure 3E**); this was not the case for MOFA+.

### Spatially-informed differential expression on mouse brain Slide-seq

We used gene expression as the primary modality, while spatial density and cell type labels were the secondary modalities. We first computed spatial density information for each cell by learning a two-dimensional Gaussian-kernel density function on cell locations; it assigns higher scores to regions with denser cell packing (**Figure 2C**). We then ran Schema using the gene expression matrix as the primary dataset, with the secondary datasets consisting of the numeric kernel-density scores, as well as the categorical labels corresponding to the four most common non-granule cell types. We aimed to find a transformation of the primary data that maximized correlation with cell spatial density while preserving a high correlation with granule cell type labels. Additionally, differences in cell-type distribution between dense and sparse regions are a confounding factor when seeking to identify a gene set specific to the granule cell type. To mitigate this, we assigned a small negative weight to correlation with non-granule cell type labels in Schema’s objective function. The primary dataset was pre-processed with a non-negative matrix factorization (NMF) transformation, limiting it to the top 100 NMF factors. Each Schema run consisted of multiple sub-runs over an ensemble of parameter settings, with the results averaged across these. The gene scores from each sub-run were a weighted average of the features with each feature’s weight as *e^w^*, *w* being the Schema-computed weights; cell loadings were computed similarly. This softmax approach is parameter-free and ensures that gene rankings are informed primarily by the features with the highest Schema weight.

To adapt CCA for a three-way modality synthesis, we tested two approaches: (1) combining spatial density and cell-type information into a composite measure that was then correlated to gene expression, or (2) performing two separate CCA analyses (correlating gene expression against either spatial density or cell type) and combining them. In the first CCA-based approach, we combined spatial density and cell-type labels by learning a Gaussian kernel density function only on cells labeled as granule cells and then inferring its value for other cells. This score was then used in CCA. In the second CCA-based approach, where we integrated results from two preliminary CCA runs, the combined cell loadings were computed as the average of the normalized cell loadings from the two CCAs, with the final genes scores then computed by a matching pursuit technique^45,46^: the final CCA score of a gene was the dot product of the CCA cell loadings and the gene’s expression vector. In our evaluations, the first CCA-based approach performed comparably or worse than the second, and the results for only the latter are presented in this paper.

We also needed to adapt SpatialDE and Trendsceek, both of which have unsupervised formulations, to select for genes whose expression shows spatial variation in granule cell types but not in non-granule cell types. To do so, we ran them separately on granule and non-granule cells and then ranked genes based on the difference of gene ranks between the two runs.

### Genomic location and accessibility inform variability in expression

We featurized chromatin accessibility by computing levels of accessibility at different locations and window sizes relative to the transcription start and end sites of a gene. For example, the feature value *rbf_20e3* for a gene was computed as follows: we first assigned weights to each ATAC-seq peak in the data, with a peak *d* base pairs upstream of the gene’s transcription start site (TSS) in the chromosome given the weight 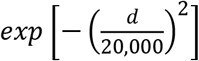 while any peak overlapping with the gene, or downstream of it, or on other chromosomes was assigned a weight of zero. The *rbf_20e3* score for the gene was the weighted sum of peak counts across all cells, with peaks in each cell weighted by the gene’s expression level. Analogously, the feature *rbf_behind_20e3* covered the region downstream of the transcription end site. We constructed multiple features, with windows of varying sizes (exponential decay factors = 20 x 10^3^, 1 x 10^6^, 10 x 10^6^ etc.) covering regions both upstream and downstream of the gene (**Figure 3B**). This feature definition avoids the need to specify arbitrary boundaries between windows. While the coverage windows of these features do overlap with each other, they differ in which gene neighborhood is emphasized in each. Our choice of the Gaussian radial basis function (*rbf*) as the weighting function was motivated by its common use in machine learning. Because of the sparsity of ATAC-seq data, features with very narrow windows were infeasible; in particular, a feature designed to cover primarily the 500 bp promoter region upstream of a gene was scored as zero too often to be informative and was removed during pre-processing. The secondary modality was constructed as follows: for each gene, we centered and *l_2_*-normalized the expression vector of the gene across all cells and sorted this vector. The Euclidean distance between two vectors then captures if there is a difference in the variability of their expression profiles. We used Schema to find a set of feature weights that maximized correlation between distances in the primary and secondary modalities. Each Schema run consisted of multiple sub-runs over an ensemble of parameter choices; the final feature weights were taken as the average across these sub-runs.

To compute feature weights using ridge regression, we used the features in the primary modality as explanatory variables and summarized each gene’s expression variability as the standard deviation of the corresponding expression vector in the secondary modality, setting this as the response variable. Schema and ridge regression both assign the highest weights to the *rbf_10e6* and *rbf_behind_10e6* features (**Figure 3C** and **Supp Table 3**). Compared to ridge regression, Schema offers the advantage of being able to correlate the primary modality with a multi-dimensional secondary modality rather than just a single summary statistic. Also, the constrained optimization framework that limits distortions of the primary modality acts as a regularization, enhancing the stability of Schema’s results. To evaluate this stability, we divided genes into subsets (by their strand orientation and chromosome); by applying Schema and ridge regression on each subset independently. We then computed the standard error in estimates of feature weight and the corresponding *t*-statistic, allowing us to compare the two approaches (**Supp Table 3**).

To test the null hypothesis that the genes in the set were dispersed randomly across TADs, we formulated two independent tests: 1) the frequency of gene pairs that co-occupy a TAD is indistinguishable from random, and 2) given the count of genes in each TAD, the rate parameter of the exponential distribution fitted to these TAD occupancy counts is indistinguishable from random. By permutation tests where we randomized gene-TAD assignments, we were able to reject the null hypothesis for the top quartile of genes (i.e. the most variable) in both cases, with Bonferroni-corrected *p* < 0.004. The trend continues across the remaining three quartiles, with the TAD clustering scores for the bottom quartile being the lowest (**Figure 3D**).

### Schema reveals CDR3 segments crucial to T-cell receptor binding specificity

When estimating location-specific selection pressure (**Figure 4C, D**), we truncated CDR3 sequences to the first 20 residues (sequences longer than that constituted less than 0.2% of our dataset). The *i*^th^ element of the primary modality feature vector was the 1-letter code of the amino acid at the *i*^th^ sequence position or a null value if the sequence length was shorter than *i*. We defined the distance between two sequences as the number of elements that disagreed. In the original space, this corresponds to the Hamming distance; in the transformed space, it is a location-weighted version of the Hamming distance. Each Schema run was an ensemble of sub-runs, with varying parameter choices of minimum correlation between the original and transformed datasets and the maximum allowed feature weights. Feature weights produced in each sub-run were normalized by linearly mapping the lowest weight to 0 and the highest to 1. We then averaged these normalized feature weights across sub-runs. To compute a location’s score using VDJdb, we extracted the VDJdb-provided relative entropy score (*I.norm*) for the location in each TCR motif and averaged it across all motifs in the database. Here, Schema and VDJdb scores have opposite orientations: for a location that demonstrates low variability, the associated Schema weight will be high while the VDJdb score will be low. Therefore, when comparing the Schema and VDJdb scores, we inverted the orientation of Schema scores by subtracting them from 1 (**Figure 4C, D**).

The comparison of per-location scores between Schema and VDJdb is complicated by length differences between motifs in VDJdb and sequences in our dataset: the former contains shorter sequences, with the average sequence length of α and β chain motifs in VDJdb being 11.9 and 12.5, respectively; the corresponding averages in our dataset are 13.5 and 14.5. However, both datasets have good coverage of locations 1-10 and the per-location scores are in broad agreement there. (**Figure 4C, D**).

To compute the selection pressure on amino acids, we focused on segments 3-7 in TCR α chains and 5-11 in TCR β chains, choosing these locations for their high sequence variability as estimated by Schema and VDJdb above. To compute Schema scores, an ensemble of sub-runs was performed, and as described above, Schema scores were normalized. VDJdb scores for an amino acid were computed as the average frequency-weighted relative entropy scores (*height.I.norm*) across the selected locations in all TCR motifs in the database.

To exemplify how Schema can synthesize additional modalities, we also incorporated proteomic measurements of 12 cell-surface markers. Hypothesizing that cell-surface protein levels should be unrelated to the V(D)J recombination variability, we added a low weight term to Schema’s objective function that *penalized* correlation between distances in the CDR3-sequence space and distances in proteomic-measurement space. Across subsets of the dataset split by donors (4 subsets) or by epitopes (10 randomly divided subsets), we compared the baseline two-modality setup against the new three-modality setup and found that the latter produced slightly more stable results than the former, with smaller standard deviations of Schema-computed weights across the subsets of data (0.094 vs 0.101 for the donor split, and 0.164 vs 0.166 for the epitope split). In general, we recommend the use of cross-validation or an independent metric to calibrate the relative weights of secondary modalities in such use-cases.

### Schema highlights secondary patterns while preserving primary structure

We chose gene expression as the primary modality, reducing it with non-negative matrix factorization (NMF) to the top 50 components, and used temporal metadata as the secondary modality. We estimated differential pseudotime using the implementation in Scanpy^36^ of Haghverdi et al.’s^59^ algorithm.

### Schema can synthesize spliced and unspliced RNA counts to accentuate cell differentiation

We normalized the spliced and unspliced counts, log transformed them and reduced them to their top 100 principal components. These were specified to Schema as the primary and secondary modalities, respectively.

To construct a pseudotime estimate from Schema’s output, we first computed the mean per-cell difference between the transformed and original RNA-seq data. Interpreting this difference as the major axis of transcriptional change, we projected the original RNA-seq values on it. The magnitude of projection for each cell is a score that we interpreted as a pseudotime measure.

## Acknowledgements

We thank B. DeMeo and the Berger laboratory members for valuable discussions and feedback. R.S. and B.H. are partially supported by the NIH grant R01 GM081871 (to B. Berger).

## Code Availability

Python source code, a tutorial, and some examples demonstrating Schema’s use are available at https://github.com/rs239/schema. The program is also available as the Python package *“schema_learn”* that can be installed using the Python package installer, pip.

## Data Availability

We used the following publicly available datasets. Below, GEO refers to the Gene Expression Omnibus repository (https://www.ncbi.nlm.nih.gov/geo/):

- Slide-seq data from Rodriques et al.^10^
  - https://singlecell.broadinstitute.org/single_cell/study/SCP354/slide-seq-study
  - Processing code at: https://github.com/broadchenf/Slideseq/tree/master/BeadSeq%20Code
- Sci-CAR (ATAC-seq and RNA-seq) data from Cao et al.^6^ (GSE117089 from GEO)
- Topologically Associating Domains data for A549 cells from ENCODE (accession ENCFF336WPU)^34,35^
- Multi-modal T-Cell receptor data from 10x Genomics^5^
  - https://www.10xgenomics.com/resources/datasets/#dataset-accordion-221-3-0-2-content
- T-cell motif data from Shugay et al’s^38^ VDJdb database:
  - Website: https://vdjdb.cdr3.net/
  - Bulk data (contains motif_pwms.txt, a file describing position-weight matrices of motifs): https://github.com/antigenomics/vdjdb-db/releases/tag/2020-01-20
  - Older version of the VDJdb motif data (does not incorporate the 10x Genomics multi-model TCR data): https://raw.githubusercontent.com/antigenomics/vdjdb-motifs/master/motif_pwms.txt
- Davie et al.’s^3^ RNA-seq data of the aging *Drosophila* brain (GSE107451 from GEO)
- Argelaguet et al.’s^21,53^ preprocessed version of Pjiuan-Sala et al.’s^57^ RNA-seq data on mouse gastrulation and their pretrained models, available at ftp://ftp.ebi.ac.uk/pub/databases/mofa/scrna_gastrulation/
- Hochgerner et al’s^58^, RNA-seq data on dentate gyrus neurogenesis (GSE95753 from GEO), made available as a sample dataset in the Python package *scvelo*.

Software: We used the following packages:

- Python (version 3.6.1): *scanpy* (version 1.5.1), *scikit-learn* (version 0.21.3), *scvelo* (version 0.2.1), *pandas* (version 0.25.1), *numpy* (version 1.17.1), *scipy* (version 1.5.1), *SpatialDE* (version 1.1.3), *leidenalg* (version 0.8.1).
- R (version 3.6.3): *MOFA2* (version 1.1), *trendsceek* (https://github.com/edsgard/trendsceek).

## Supplementary Materials

**Supp Figure 1.**
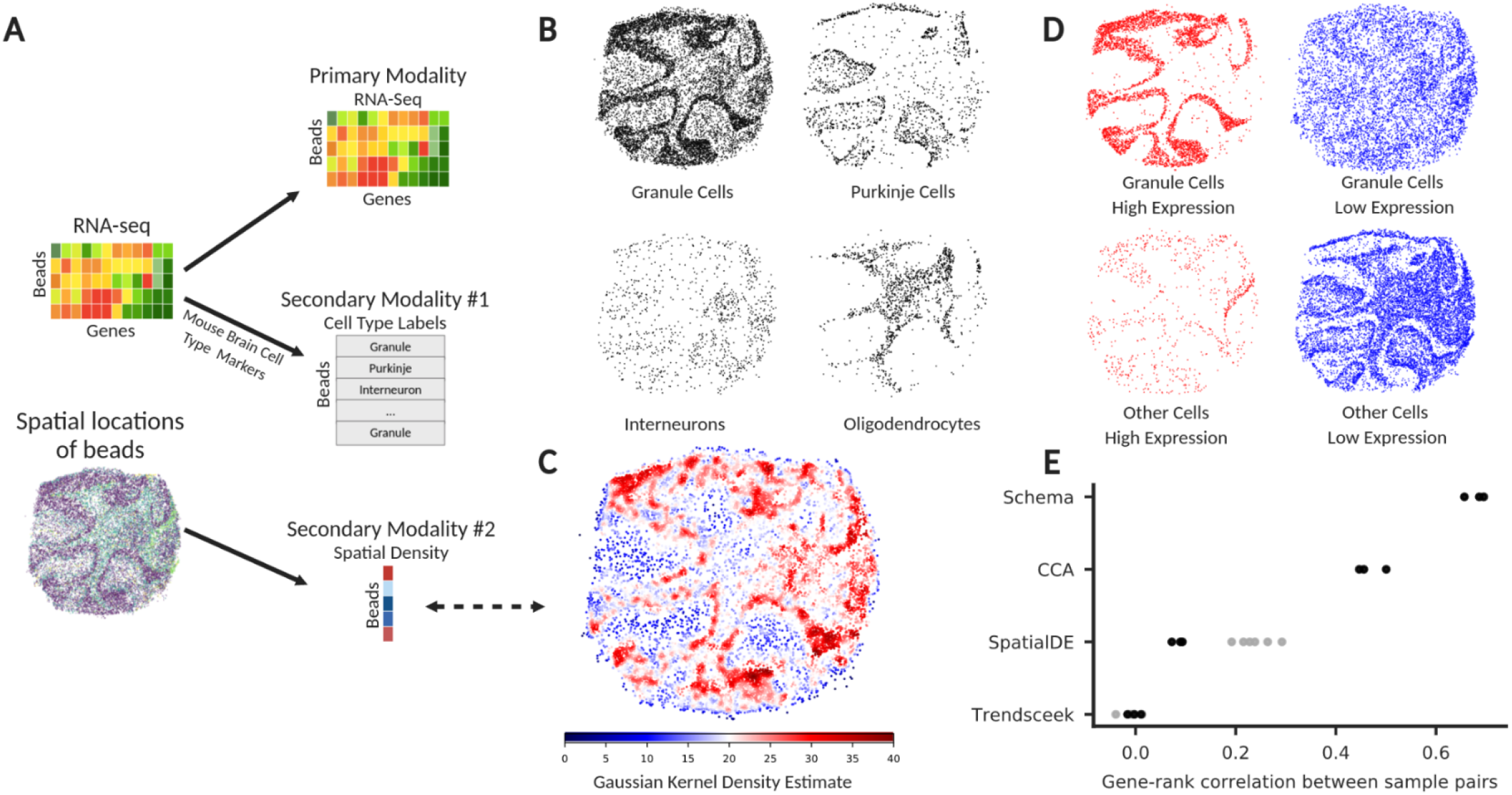
Schema identifies a gene set in granule neurons whose expression covaries with spatial cellular density. (**A**) Rodriques et al.^10^ simultaneously assayed spatial and transcriptomic modalities in mouse cerebellum tissue (here, data from puck 180430_1 is shown). In addition, they labeled beads (each corresponding to a transcriptome) with putative cell-type by comparing gene expression profiles with known cell-type markers. (**B**) Spatial distribution of the most common cell types in the tissue section: granule cells, Purkinje cells, interneurons, and oligodendrocytes. Note the variation in spatial density for granule cells. (**C**) We quantified this spatial density variation by computing a two-dimensional Gaussian-kernel density estimate, with cells in dense regions assigned a higher score. (**D**) Schema is able to identify a ranked set of genes that are highly expressed only in densely-packed granule cells. The four figures here show mutually disjoint sets of cells: granule cells with high expression of the gene set, granule cells with low expression of the gene set, other cells with high expression, and other cells with low expression. Here, a cell is said to have high expression of the gene set if the cell’s loading on this gene set ranks in the top quartile. (**E**) Evaluation of the stability of gene rankings computed by Schema, canonical correlation analysis (CCA), SpatialDE and Trendsceek on three replicates sourced from mouse cerebellum tissue. The black points indicate the Spearman rank correlation of gene scores across pairs of replicates. To adapt SpatialDE and Trendsceek for this task, we first ran them separately on granule and non-granule cells, scoring genes by spatial variability in expression; the final result (black points) was then computed as a rank-difference of the cell-type specific scores. Here, the grey points show the cross-replicate gene-rank correlation of the intermediate results.

**Supp Figure 2.**
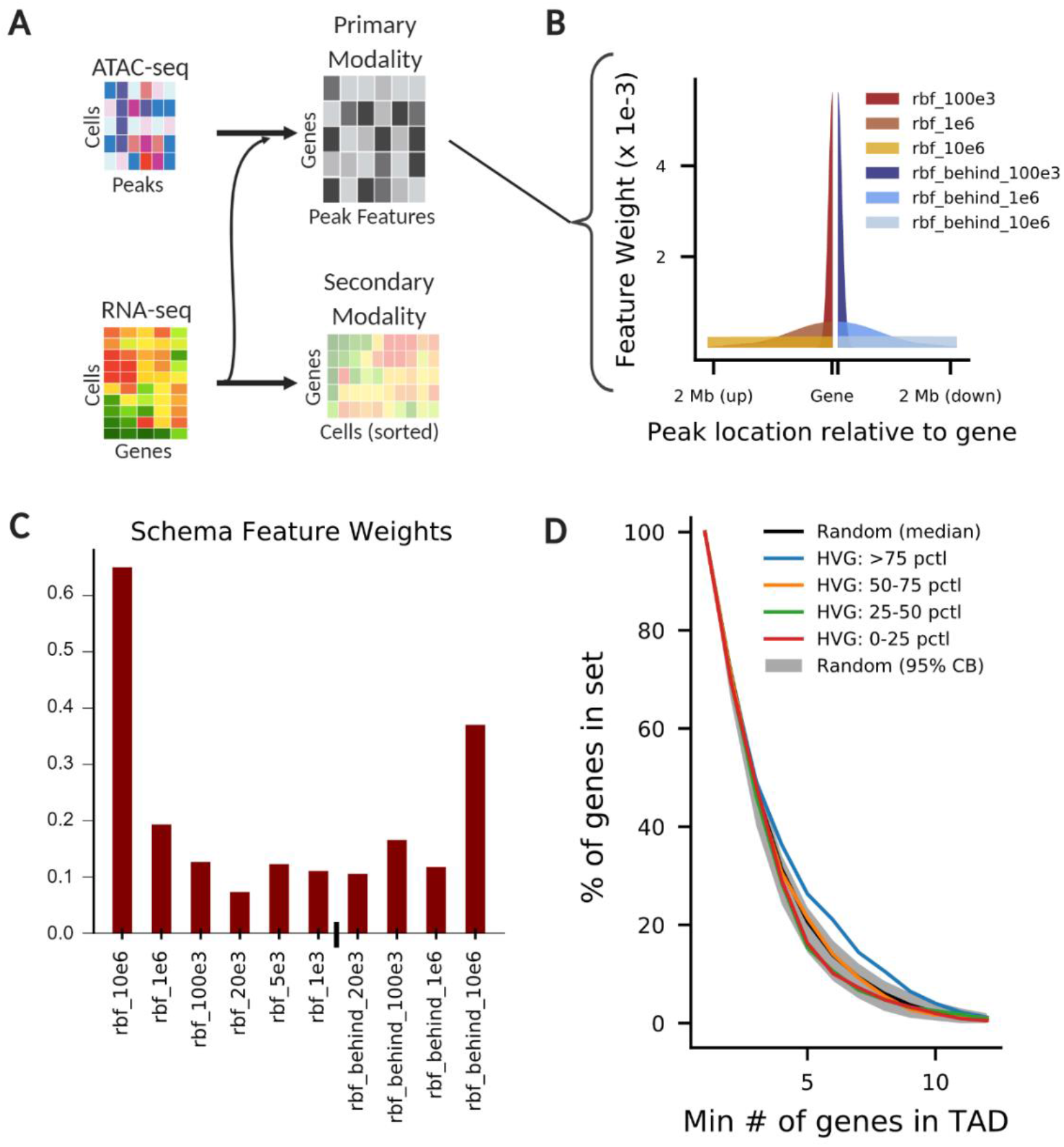
Highly variable genes are significantly more likely to cluster together in topologically associating domains (TADs) (**A**) We investigated expression variability in the context of chromatin accessibility, evaluating if highly variable genes display differential accessibility around their genomic loci. With genes as the units of observation, we used Schema to analyze simultaneously assayed ATAC-seq and RNA-seq data^6^. The primary modality captured peak counts around a gene while the distance metric on the secondary modality captured if two genes have similar levels of expression variability. (**B**) Each feature of the primary modality corresponded to a genomic region near the gene, scoring how its expression covaries with peak counts in the region. The feature’s range is defined by a Gaussian radial basis function, of the form *exp*(–(*d* ≥ *λ*)^2), that weights the contribution of a peak by its distance *d* from the gene. We defined features upstream and downstream of the gene’s transcription start and end sites, respectively. (**C**) Schema identifies long-range features (~10 Mb) as being the most relevant in correlating chromatin accessibility with expression variability. (**D**) To further explore this result, we investigated the organization of highly variable genes in topologically associating domains (TADs). We divided genes into quartiles by expression variability. For each quartile, we plotted the fraction of genes that are in TADs containing *k* or more genes from the set (*k* is on the *x*-axis). The gray region and the black line represent the 95% range and median, respectively, of random baselines generated by shuffling genes between TADs. Genes in the top quartile by expression variability (blue curve) are significantly likely to be clustered together into TADs (Bonferroni-corrected *ρ* < 0.004, permutation test)

**Supp Figure 3.**
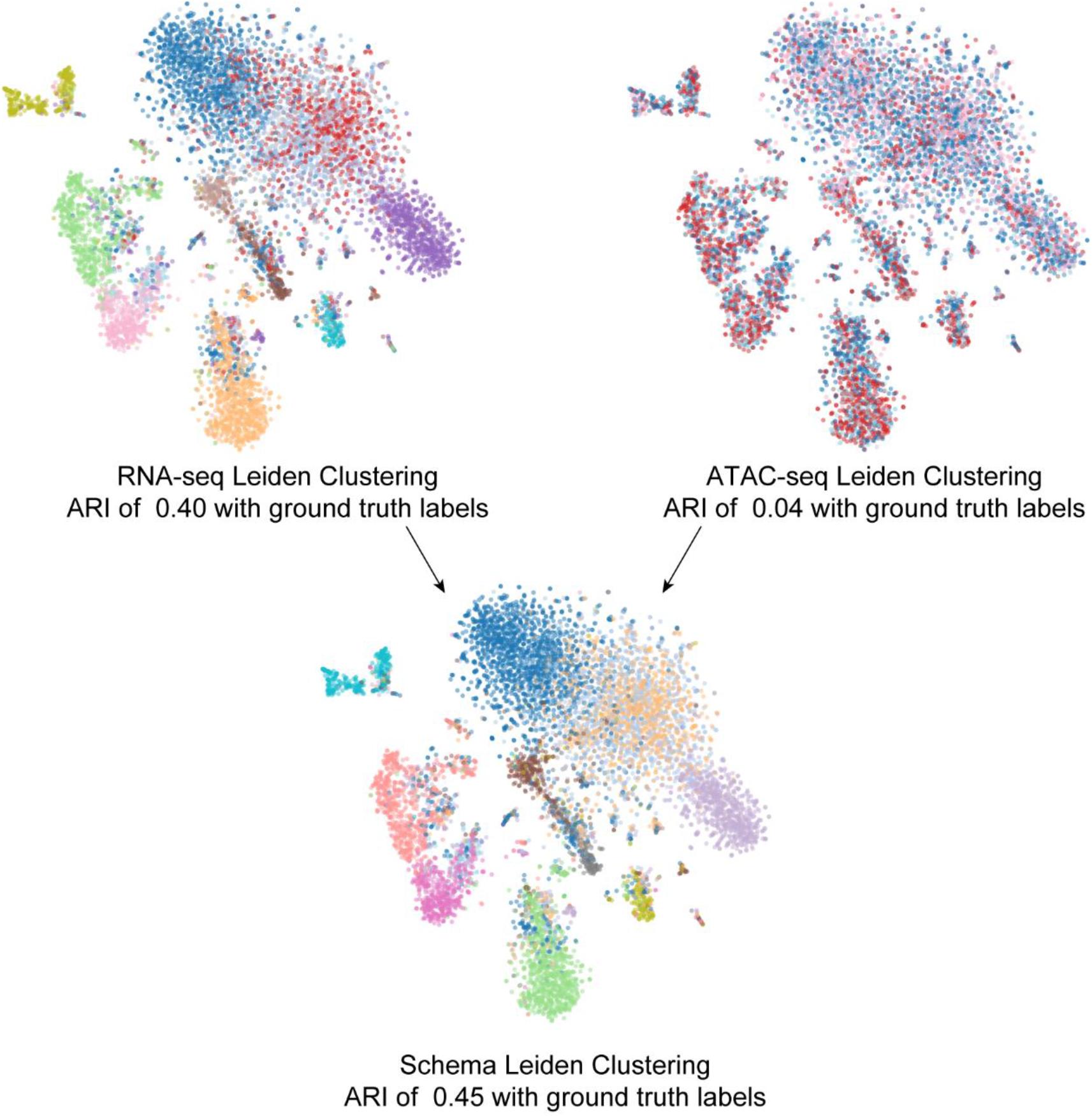
Leiden clustering (and its ARI against ground truth) for RNA-seq and ATAC-seq data individually

**Supp Figure 4.**
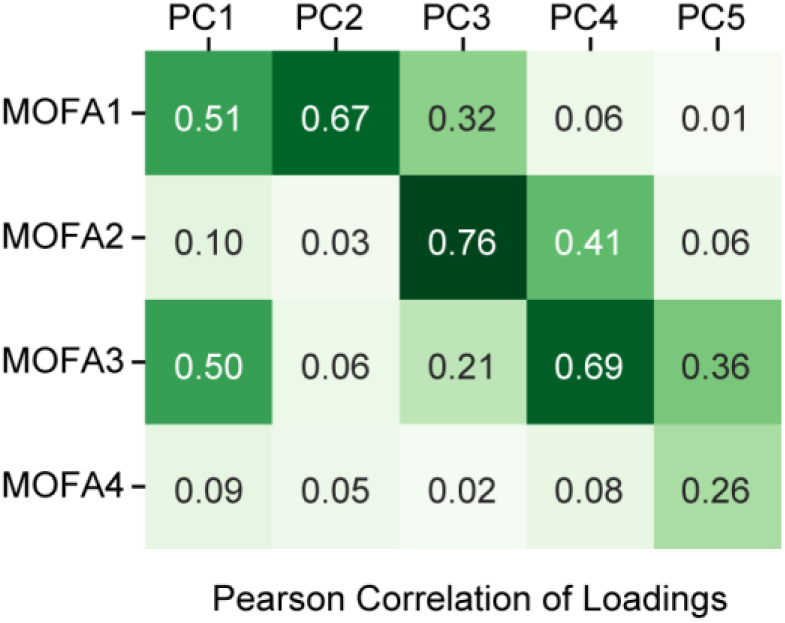
Correlation of factor loadings between MOFA+ factors and principal components

**Supp Table 1.** REACTOME^27^ pathways enriched in Schema-ranked genes across 3 samples of Slide-Seq data

| Pathway identifier | Pathway name | Entities FDR |
| --- | --- | --- |
| R-HSA-156902 | Peptide chain elongation | 6.44E-15 |
| R-HSA-72689 | Formation of a pool of free 40S subunits | 6.44E-15 |
| R-HSA-975956 | Nonsense Mediated Decay (NMD) independent of the Exon Junction Complex (EJC) | 6.44E-15 |
| R-HSA-72706 | GTP hydrolysis and joining of the 60S ribosomal subunit | 6.44E-15 |
| R-HSA-156827 | L13a-mediated translational silencing of Ceruloplasmin expression | 6.44E-15 |
| R-HSA-927802 | Nonsense-Mediated Decay (NMD) | 6.44E-15 |
| R-HSA-975957 | Nonsense Mediated Decay (NMD) enhanced by the Exon Junction Complex (EJC) | 6.44E-15 |
| R-HSA-1799339 | SRP-dependent cotranslational protein targeting to membrane | 6.44E-15 |
| R-HSA-192823 | Viral mRNA Translation | 6.44E-15 |
| R-HSA-72613 | Eukaryotic Translation Initiation | 6.44E-15 |
| R-HSA-72737 | Cap-dependent Translation Initiation | 6.44E-15 |
| R-HSA-6799198 | Complex I biogenesis | 6.44E-15 |
| R-HSA-156842 | Eukaryotic Translation Elongation | 6.44E-15 |
| R-HSA-611105 | Respiratory electron transport | 6.44E-15 |
| R-HSA-163200 | Respiratory electron transport, ATP synthesis by chemiosmotic coupling, and heat production by uncoupling proteins. | 6.44E-15 |
| R-HSA-72766 | Translation | 6.44E-15 |
| R-HSA-72764 | Eukaryotic Translation Termination | 6.44E-15 |
| R-HSA-8953854 | Metabolism of RNA | 6.44E-15 |
| R-HSA-1428517 | The citric acid (TCA) cycle and respiratory electron transport | 6.44E-15 |
| R-HSA-2262752 | Cellular responses to stress | 6.44E-15 |
| R-HSA-8953897 | Cellular responses to external stimuli | 6.44E-15 |
| R-HSA-422475 | Axon guidance | 6.44E-15 |
| R-HSA-168273 | Influenza Viral RNA Transcription and Replication | 6.44E-15 |
| R-HSA-9633012 | Response of EIF2AK4 (GCN2) to amino acid deficiency | 6.44E-15 |
| R-HSA-9010553 | Regulation of expression of SLITs and ROBOs | 6.44E-15 |
| R-HSA-2408557 | Selenocysteine synthesis | 6.44E-15 |
| R-HSA-376176 | Signaling by ROBO receptors | 6.44E-15 |
| R-HSA-168254 | Influenza Infection | 6.44E-15 |
| R-HSA-168255 | Influenza Life Cycle | 6.44E-15 |
| R-HSA-6791226 | Major pathway of rRNA processing in the nucleolus and cytosol | 2.49E-14 |
| R-HSA-72649 | Translation initiation complex formation | 1.20E-13 |
| R-HSA-2408522 | Selenoamino acid metabolism | 1.67E-13 |
| R-HSA-72312 | rRNA processing | 2.72E-13 |
| R-HSA-72702 | Ribosomal scanning and start codon recognition | 2.72E-13 |
| R-HSA-8868773 | rRNA processing in the nucleus and cytosol | 3.84E-13 |
| R-HSA-72662 | Activation of the mRNA upon binding of the cap-binding complex and eIFs, and subsequent binding to 43S | 6.03E-13 |
| R-HSA-72695 | Formation of the ternary complex, and subsequently, the 43S complex | 3.17E-12 |
| R-HSA-5663205 | Infectious disease | 2.12E-11 |
| R-HSA-9604323 | Negative regulation of NOTCH4 signaling | 9.07E-09 |
| R-HSA-72163 | mRNA Splicing - Major Pathway | 1.12E-08 |
| R-HSA-69275 | G2/M Transition | 1.7E-08 |
| R-HSA-453274 | Mitotic G2-G2/M phases | 2.29E-08 |
| R-HSA-195253 | Degradation of beta-catenin by the destruction complex | 4.37E-08 |
| R-HSA-5610785 | GLI3 is processed to GLI3R by the proteasome | 4.41E-08 |
| R-HSA-5610783 | Degradation of GLI2 by the proteasome | 4.41E-08 |
| R-HSA-72172 | mRNA Splicing | 6.28E-08 |
| R-HSA-3858494 | Beta-catenin independent WNT signaling | 6.59E-08 |
| R-HSA-8852276 | The role of GTSE1 in G2/M progression after G2 checkpoint | 6.94E-08 |
| R-HSA-8854050 | FBXL7 down-regulates AURKA during mitotic entry and in early mitosis | 8.16E-08 |
| R-HSA-450408 | AUF1 (hnRNP D0) binds and destabilizes mRNA | 1.12E-07 |
| R-HSA-180585 | Vif-mediated degradation of APOBEC3G | 1.12E-07 |
| R-HSA-1234176 | Oxygen-dependent proline hydroxylation of Hypoxia-inducible Factor Alpha | 1.89E-07 |
| R-HSA-1266738 | Developmental Biology | 2.40E-07 |
| R-HSA-162909 | Host Interactions of HIV factors | 2.40E-07 |
| R-HSA-4086400 | PCP/CE pathway | 2.40E-07 |
| R-HSA-392499 | Metabolism of proteins | 3.30E-07 |
| R-HSA-5610787 | Hedgehog 'off' state | 4.17E-07 |
| R-HSA-163210 | Formation of ATP by chemiosmotic coupling | 4.22E-07 |
| R-HSA-72203 | Processing of Capped Intron-Containing Pre-mRNA | 4.22E-07 |
| R-HSA-8949613 | Cristae formation | 5.75E-07 |
| R-HSA-187577 | SCF(Skp2)-mediated degradation of p27/p21 | 5.86E-07 |
| R-HSA-5610780 | Degradation of GLI1 by the proteasome | 5.86E-07 |
| R-HSA-450531 | Regulation of mRNA stability by proteins that bind AU-rich elements | 6.28E-07 |
| R-HSA-180534 | Vpu mediated degradation of CD4 | 6.50E-07 |
| R-HSA-195721 | Signaling by WNT | 8.67E-07 |
| R-HSA-75815 | Ubiquitin-dependent degradation of Cyclin D | 8.95E-07 |
| R-HSA-199991 | Membrane Trafficking | 9.16E-07 |
| R-HSA-211733 | Regulation of activated PAK-2p34 by proteasome mediated degradation | 1.056E-06 |
| R-HSA-1234174 | Cellular response to hypoxia | 1.10E-06 |
| R-HSA-174113 | SCF-beta-TrCP mediated degradation of Emi1 | 1.13E-06 |
| R-HSA-112316 | Neuronal System | 1.29E-06 |
| R-HSA-5658442 | Regulation of RAS by GAPs | 1.52E-06 |
| R-HSA-5362768 | Hh mutants that don't undergo autocatalytic processing are degraded by ERAD | 1.53E-06 |
| R-HSA-1168372 | Downstream signaling events of B Cell Receptor (BCR) | 1.82E-06 |
| R-HSA-4641258 | Degradation of DVL | 1.884E-06 |
| R-HSA-8939902 | Regulation of RUNX2 expression and activity | 2.11E-06 |
| R-HSA-174084 | Autodegradation of Cdh1 by Cdh1:APC/C | 2.56E-06 |
| R-HSA-373760 | L1CAM interactions | 2.81E-06 |
| R-HSA-69613 | p53-Independent G1/S DNA damage checkpoint | 2.93E-06 |
| R-HSA-69601 | Ubiquitin Mediated Degradation of Phosphorylated Cdc25A | 2.93E-06 |
| R-HSA-69610 | p53-Independent DNA Damage Response | 2.93E-06 |
| R-HSA-349425 | Autodegradation of the E3 ubiquitin ligase COP1 | 2.93E-06 |
| R-HSA-169911 | Regulation of Apoptosis | 2.93E-06 |
| R-HSA-5387390 | Hh mutants abrogate ligand secretion | 3.04E-06 |
| R-HSA-162906 | HIV Infection | 3.29E-06 |
| R-HSA-77387 | Insulin receptor recycling | 3.48E-06 |
| R-HSA-5676590 | NIK-->noncanonical NF-kB signaling | 5.00E-06 |
| R-HSA-69202 | Cyclin E associated events during G1/S transition | 5.58E-06 |
| R-HSA-174178 | APC/C:Cdh1 mediated degradation of Cdc20 and other APC/C:Cdh1 targeted proteins in late mitosis/early G1 | 5.93E-06 |
| R-HSA-174154 | APC/C:Cdc20 mediated degradation of Securin | 6.231E-06 |
| R-HSA-5678895 | Defective CFTR causes cystic fibrosis | 6.231E-06 |
| R-HSA-4641257 | Degradation of AXIN | 6.231E-06 |
| R-HSA-8941858 | Regulation of RUNX3 expression and activity | 6.231E-06 |
| R-HSA-5358351 | Signaling by Hedgehog | 6.89E-06 |
| R-HSA-68949 | Orc1 removal from chromatin | 7.13E-06 |
| R-HSA-69656 | Cyclin A:Cdk2-associated events at S phase entry | 8.06E-06 |
| R-HSA-112315 | Transmission across Chemical Synapses | 9.99E-06 |
| R-HSA-68827 | CDT1 association with the CDC6:ORC:origin complex | 1.01E-05 |
| R-HSA-69541 | Stabilization of p53 | 1.01E-05 |
| R-HSA-9013694 | Signaling by NOTCH4 | 1.15E-05 |
| R-HSA-69563 | p53-Dependent G1 DNA Damage Response | 1.17E-05 |
| R-HSA-69580 | p53-Dependent G1/S DNA damage checkpoint | 1.17E-05 |
| R-HSA-195258 | RHO GTPase Effectors | 1.18E-05 |
| R-HSA-4608870 | Asymmetric localization of PCP proteins | 1.50E-05 |
| R-HSA-5607761 | Dectin-1 mediated noncanonical NF-kB signaling | 1.50E-05 |
| R-HSA-350562 | Regulation of ornithine decarboxylase (ODC) | 1.63E-05 |
| R-HSA-5687128 | MAPK6/MAPK4 signaling | 1.63E-05 |
| R-HSA-917937 | Iron uptake and transport | 1.63E-05 |
| R-HSA-174184 | Cdc20:Phospho-APC/C mediated degradation of Cyclin A | 1.63E-05 |
| R-HSA-1169091 | Activation of NF-kappaB in B cells | 1.63E-05 |
| R-HSA-69615 | G1/S DNA Damage Checkpoints | 1.63E-05 |
| R-HSA-5358346 | Hedgehog ligand biogenesis | 1.63E-05 |
| R-HSA-179419 | APC:Cdc20 mediated degradation of cell cycle proteins prior to satisfaction of the cell cycle checkpoint | 2.05E-05 |
| R-HSA-112314 | Neurotransmitter receptors and postsynaptic signal transmission | 2.14E-05 |
| R-HSA-8948751 | Regulation of PTEN stability and activity | 2.56E-05 |
| R-HSA-69017 | CDK-mediated phosphorylation and removal of Cdc6 | 2.56E-05 |
| R-HSA-69052 | Switching of origins to a post-replicative state | 2.90E-05 |
| R-HSA-176409 | APC/C:Cdc20 mediated degradation of mitotic proteins | 2.9553E-05 |
| R-HSA-176814 | Activation of APC/C and APC/C:Cdc20 mediated degradation of mitotic proteins | 3.66E-05 |
| R-HSA-109581 | Apoptosis | 3.71E-05 |
| R-HSA-176408 | Regulation of APC/C activators between G1/S and early anaphase | 3.93E-05 |
| R-HSA-453276 | Regulation of mitotic cell cycle | 6.03E-05 |
| R-HSA-174143 | APC/C-mediated degradation of cell cycle proteins | 6.03E-05 |
| R-HSA-2467813 | Separation of Sister Chromatids | 6.42E-05 |
| R-HSA-68882 | Mitotic Anaphase | 6.42E-05 |
| R-HSA-68867 | Assembly of the pre-replicative complex | 6.6435E-05 |
| R-HSA-2132295 | MHC class II antigen presentation | 7.33E-05 |
| R-HSA-194315 | Signaling by Rho GTPases | 7.57E-05 |
| R-HSA-5632684 | Hedgehog 'on' state | 8.05E-05 |
| R-HSA-1236978 | Cross-presentation of soluble exogenous antigens (endosomes) | 8.72E-05 |
| R-HSA-5357801 | Programmed Cell Death | 9.0068E-05 |
| R-HSA-2555396 | Mitotic Metaphase and Anaphase | 9.03E-05 |
| R-HSA-389957 | Prefoldin mediated transfer of substrate to CCT/TriC | 9.48E-05 |
| R-HSA-3371497 | HSP90 chaperone cycle for steroid hormone receptors (SHR) | 0.00010202 |
| R-HSA-917977 | Transferrin endocytosis and recycling | 0.0001193 |
| R-HSA-5607764 | CLEC7A (Dectin-1) signaling | 0.00012677 |
| R-HSA-9020702 | Interleukin-1 signaling | 0.00016409 |
| R-HSA-8856688 | Golgi-to-ER retrograde transport | 0.00016462 |
| R-HSA-8950505 | Gene and protein expression by JAK-STAT signaling after Interleukin-12 stimulation | 0.00017241 |
| R-HSA-983168 | Antigen processing: Ubiquitination & Proteasome degradation | 0.00019434 |
| R-HSA-5689603 | UCH proteinases | 0.00020826 |
| R-HSA-2565942 | Regulation of PLK1 Activity at G2/M Transition | 0.00021349 |
| R-HSA-389958 | Cooperation of Prefoldin and TriC/CCT in actin and tubulin folding | 0.00025458 |
| R-HSA-5628897 | TP53 Regulates Metabolic Genes | 0.00026315 |
| R-HSA-391251 | Protein folding | 0.00027505 |
| R-HSA-8939236 | RUNX1 regulates transcription of genes involved in differentiation of HSCs | 0.00027505 |
| R-HSA-390466 | Chaperonin-mediated protein folding | 0.00028809 |
| R-HSA-9612973 | Autophagy | 0.00031906 |
| R-HSA-9663891 | Selective autophagy | 0.0003353 |
| R-HSA-9020591 | Interleukin-12 signaling | 0.00039889 |
| R-HSA-9639288 | Amino acids regulate mTORC1 | 0.00044699 |
| R-HSA-5653656 | Vesicle-mediated transport | 0.00051066 |
| R-HSA-68886 | M Phase | 0.00052396 |
| R-HSA-111885 | Opioid Signalling | 0.00052761 |
| R-HSA-5619084 | ABC transporter disorders | 0.00060031 |
| R-HSA-6807878 | COPI-mediated anterograde transport | 0.00075601 |
| R-HSA-69002 | DNA Replication Pre-Initiation | 0.00077512 |
| R-HSA-9646399 | Aggrephagy | 0.00080517 |
| R-HSA-2682334 | EPH-Ephrin signaling | 0.00081033 |
| R-HSA-380284 | Loss of proteins required for interphase microtubule organization from the centrosome | 0.00098495 |
| R-HSA-380259 | Loss of Nlp from mitotic centrosomes | 0.00098495 |
| R-HSA-447115 | Interleukin-12 family signaling | 0.00098495 |
| R-HSA-416993 | Trafficking of GluR2-containing AMPA receptors | 0.00098495 |

**Supp Figure 5.**
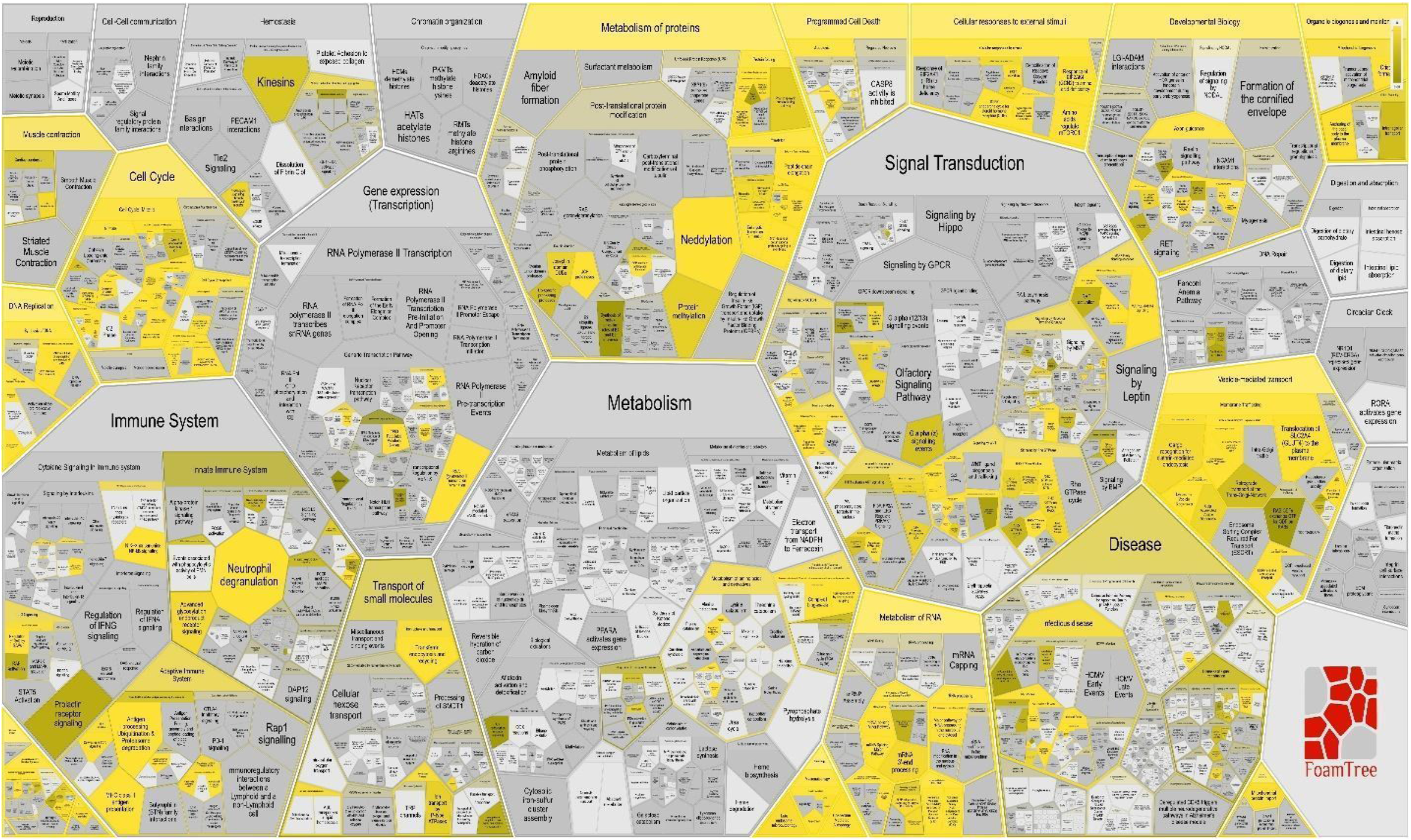
Voronoi-tessellation visualization of REACTOME pathways enriched in Schema-ranked genes across 3 samples of Slide-Seq data

**Supp Table 2.** **Gene Ontology (Biological Function) enrichment analysis of Schema-ranked genes across 3 samples of Slide-Seq data** (via GOrilla^28^)

| GO Term | Description | FDR q-value |
| --- | --- | --- |
| GO:0003735 | structural constituent of ribosome | 1.99E-24 |
| GO:0003723 | RNA binding | 7.42E-20 |
| GO:0015078 | proton transmembrane transporter activity | 2.06E-19 |
| GO:0022853 | active ion transmembrane transporter activity | 1.80E-17 |
| GO:0042625 | ATPase coupled ion transmembrane transporter activity | 1.44E-17 |
| GO:0019829 | cation-transporting ATPase activity | 1.20E-17 |
| GO:0044769 | ATPase activity, coupled to transmembrane movement of ions, rotational mechanism | 6.27E-16 |
| GO:0005198 | structural molecule activity | 1.73E-14 |
| GO:0003729 | mRNA binding | 1.67E-14 |
| GO:0015077 | monovalent inorganic cation transmembrane transporter activity | 7.49E-14 |
| GO:0003954 | NADH dehydrogenase activity | 6.29E-13 |
| GO:0008137 | NADH dehydrogenase (ubiquinone) activity | 1.73E-12 |
| GO:0050136 | NADH dehydrogenase (quinone) activity | 1.59E-12 |
| GO:0044877 | protein-containing complex binding | 1.82E-12 |
| GO:0043492 | ATPase activity, coupled to movement of substances | 7.04E-12 |
| GO:0042626 | ATPase activity, coupled to transmembrane movement of substances | 6.60E-12 |
| GO:0046933 | proton-transporting ATP synthase activity, rotational mechanism | 8.56E-12 |
| GO:0005515 | protein binding | 1.00E-11 |
| GO:0015399 | primary active transmembrane transporter activity | 1.83E-11 |
| GO:0015405 | P-P-bond-hydrolysis-driven transmembrane transporter activity | 1.74E-11 |
| GO:0019899 | enzyme binding | 2.72E-11 |
| GO:0004129 | cytochrome-c oxidase activity | 2.98E-11 |
| GO:0016676 | oxidoreductase activity, acting on a heme group of donors, oxygen as acceptor | 2.85E-11 |
| GO:0015002 | heme-copper terminal oxidase activity | 2.73E-11 |
| GO:0009055 | electron transfer activity | 2.87E-11 |
| GO:0016675 | oxidoreductase activity, acting on a heme group of donors | 5.24E-11 |
| GO:0022890 | inorganic cation transmembrane transporter activity | 2.85E-10 |
| GO:0016655 | oxidoreductase activity, acting on NAD(P)H, quinone or similar compound as acceptor | 2.93E-10 |
| GO:0005488 | binding | 8.00E-10 |
| GO:0017111 | nucleoside-triphosphatase activity | 3.32E-09 |
| GO:0019843 | rRNA binding | 3.87E-09 |
| GO:0008324 | cation transmembrane transporter activity | 6.78E-09 |
| GO:0036442 | proton-exporting ATPase activity | 1.13E-08 |
| GO:0016462 | pyrophosphatase activity | 2.62E-08 |
| GO:0051082 | unfolded protein binding | 3.08E-08 |
| GO:0016817 | hydrolase activity, acting on acid anhydrides | 3.08E-08 |
| GO:0016818 | hydrolase activity, acting on acid anhydrides, in phosphorus-containing anhydrides | 3E-08 |
| GO:0019904 | protein domain specific binding | 5.02E-08 |
| GO:0015318 | inorganic molecular entity transmembrane transporter activity | 7E-08 |
| GO:0042623 | ATPase activity, coupled | 1.28E-07 |
| GO:0015662 | ATPase activity, coupled to transmembrane movement of ions, phosphorylative mechanism | 1.65E-07 |
| GO:0016679 | oxidoreductase activity, acting on diphenols and related substances as donors | 2.25E-07 |
| GO:0016681 | oxidoreductase activity, acting on diphenols and related substances as donors, cytochrome as acceptor | 2.2E-07 |
| GO:0008121 | ubiquinol-cytochrome-c reductase activity | 2.15E-07 |
| GO:0015075 | ion transmembrane transporter activity | 3.05E-07 |
| GO:0016887 | ATPase activity | 3.46E-07 |
| GO:0031625 | ubiquitin protein ligase binding | 6.31E-07 |
| GO:0008092 | cytoskeletal protein binding | 8.32E-07 |
| GO:0044325 | ion channel binding | 2.34E-06 |
| GO:0030235 | nitric-oxide synthase regulator activity | 3.54E-06 |
| GO:0022857 | transmembrane transporter activity | 3.72E-06 |
| GO:0044389 | ubiquitin-like protein ligase binding | 4.49E-06 |
| GO:0017075 | syntaxin-1 binding | 5.79E-06 |
| GO:1990446 | U1 snRNP binding | 0.0000057 |
| GO:0005215 | transporter activity | 6.27E-06 |
| GO:0004298 | threonine-type endopeptidase activity | 6.45E-06 |
| GO:0070003 | threonine-type peptidase activity | 6.34E-06 |
| GO:0046961 | proton-transporting ATPase activity, rotational mechanism | 6.51E-06 |
| GO:0043021 | ribonucleoprotein complex binding | 0.0000065 |
| GO:0008553 | proton-exporting ATPase activity, phosphorylative mechanism | 0.0000152 |
| GO:0048306 | calcium-dependent protein binding | 0.0000223 |
| GO:0031800 | type 3 metabotropic glutamate receptor binding | 0.0000257 |
| GO:0008135 | translation factor activity, RNA binding | 0.000037 |
| GO:0016651 | oxidoreductase activity, acting on NAD(P)H | 0.0000534 |
| GO:0031386 | protein tag | 0.0000651 |
| GO:0022804 | active transmembrane transporter activity | 0.0000643 |
| GO:0005200 | structural constituent of cytoskeleton | 0.0000856 |
| GO:0015631 | tubulin binding | 0.000089 |
| GO:0019905 | syntaxin binding | 0.0000974 |
| GO:0003743 | translation initiation factor activity | 0.00011 |
| GO:0016491 | oxidoreductase activity | 0.000112 |
| GO:0031072 | heat shock protein binding | 0.000113 |
| GO:0070990 | snRNP binding | 0.00018 |
| GO:0000149 | SNARE binding | 0.000209 |
| GO:0051087 | chaperone binding | 0.000222 |
| GO:0050998 | nitric-oxide synthase binding | 0.000316 |
| GO:0003676 | nucleic acid binding | 0.000444 |
| GO:0008556 | potassium-transporting ATPase activity | 0.000576 |
| GO:0005391 | sodium:potassium-exchanging ATPase activity | 0.000568 |
| GO:1901363 | heterocyclic compound binding | 0.000706 |
| GO:0035256 | G protein-coupled glutamate receptor binding | 0.000811 |
| GO:0043532 | angiostatin binding | 0.000806 |
| GO:0003924 | GTPase activity | 0.000852 |
| GO:0097159 | organic cyclic compound binding | 0.00131 |
| GO:0047485 | protein N-terminus binding | 0.00172 |
| GO:0008179 | adenylate cyclase binding | 0.00173 |
| GO:0002135 | CTP binding | 0.00175 |
| GO:0019900 | kinase binding | 0.00214 |
| GO:0001883 | purine nucleoside binding | 0.00244 |
| GO:0003730 | mRNA 3'-UTR binding | 0.00359 |
| GO:0048156 | tau protein binding | 0.00415 |
| GO:0019901 | protein kinase binding | 0.00425 |

**Supp Figure 6.**
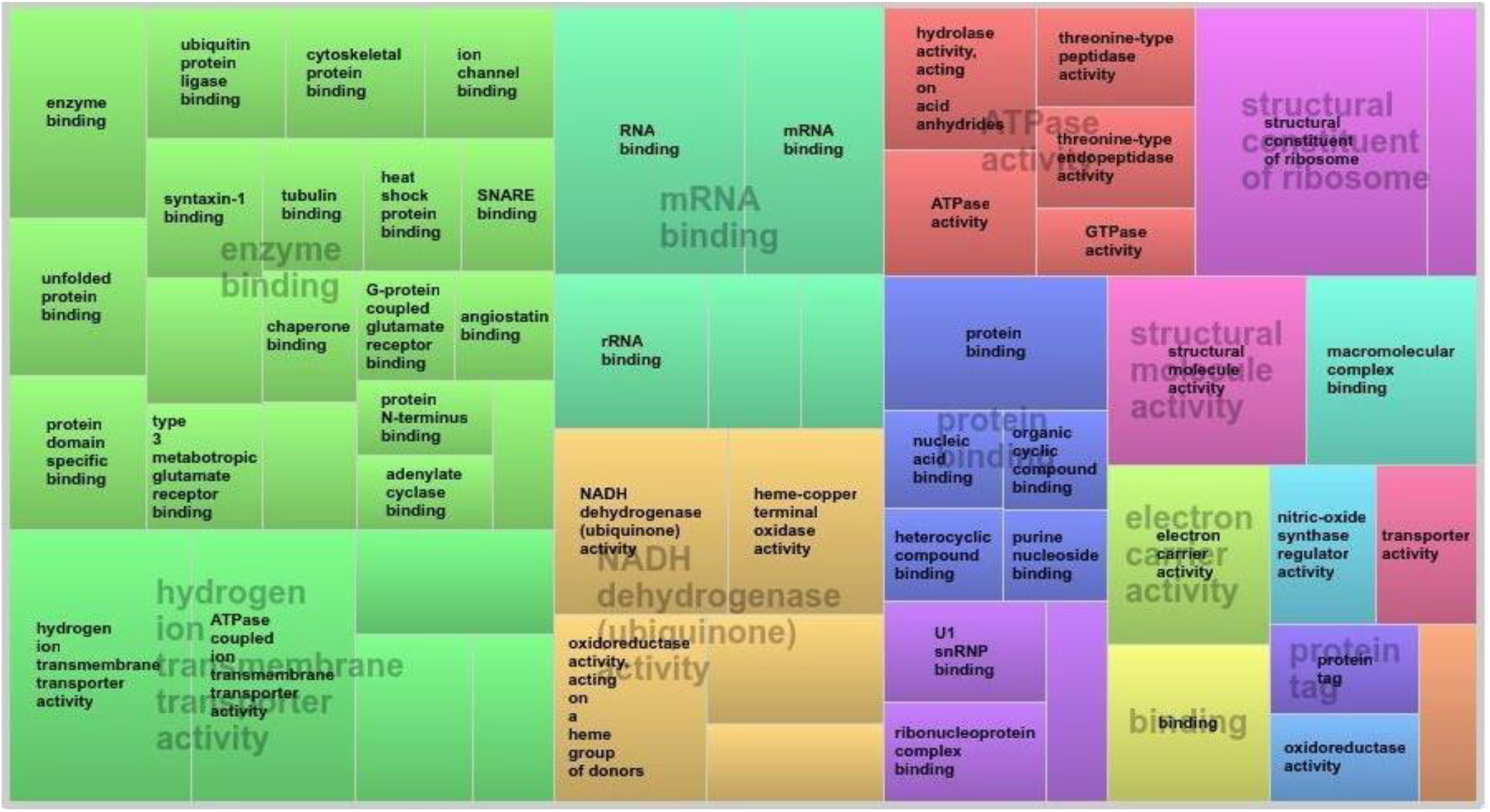
**Visualization of enriched GO terms in Schema-ranked genes across 3 samples of Slide-Seq data** (via REViGO^29^, using abs(log10(*p*-value)))

**Supp Figure 7.**
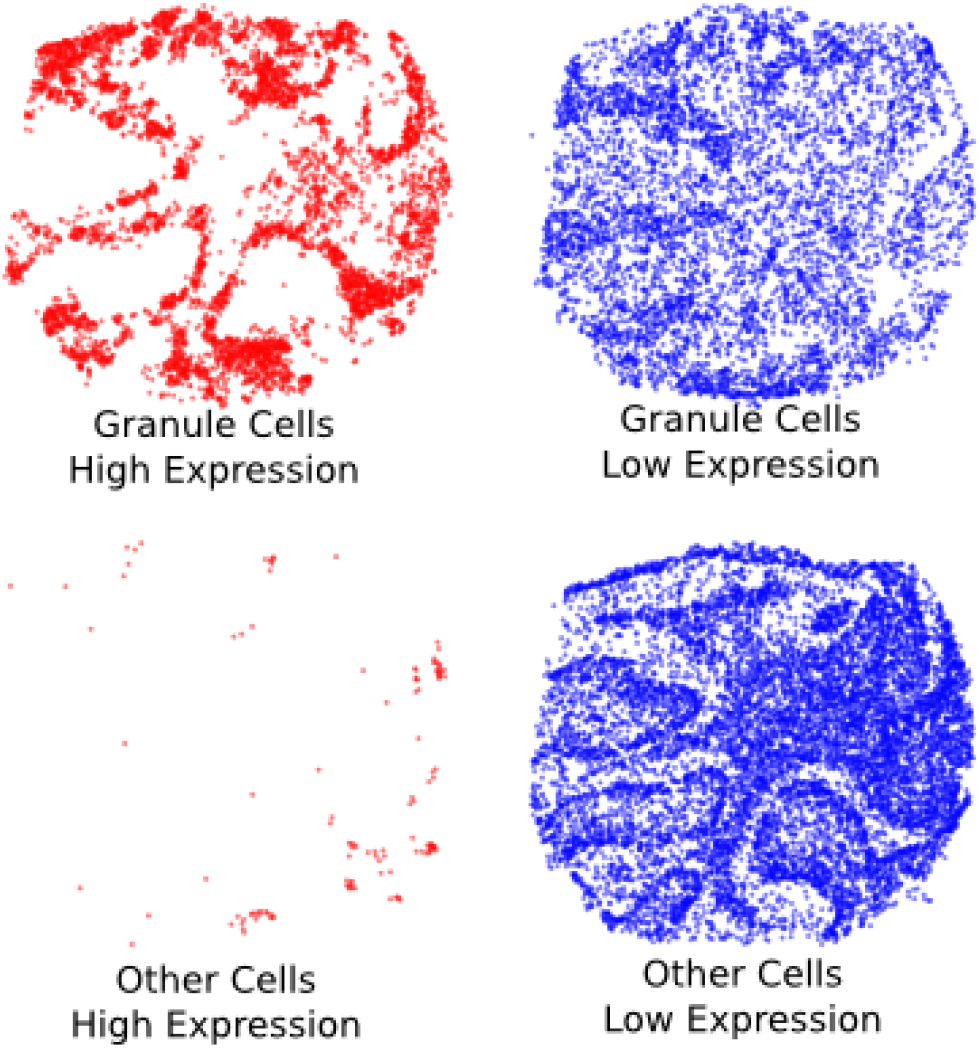
Comparison of Schema with CCA on Slide-seq data (Part 1) Visually, CCA seems to be effective at identifying a gene set that is differentially expressed only in densely-located granule neurons. The Slide-seq sample used here (Puck 180430_1) is the same as in **Supp Figure 1D**. However, as shown in **Supp Figure 2E**, the gene ranking computed by Schema is better preserved across three Slide-Seq samples than those produced by CCA (median sample-pair Spearman rank correlation of 0.675 and 0.457, respectively)

**Supp Figure 8.**
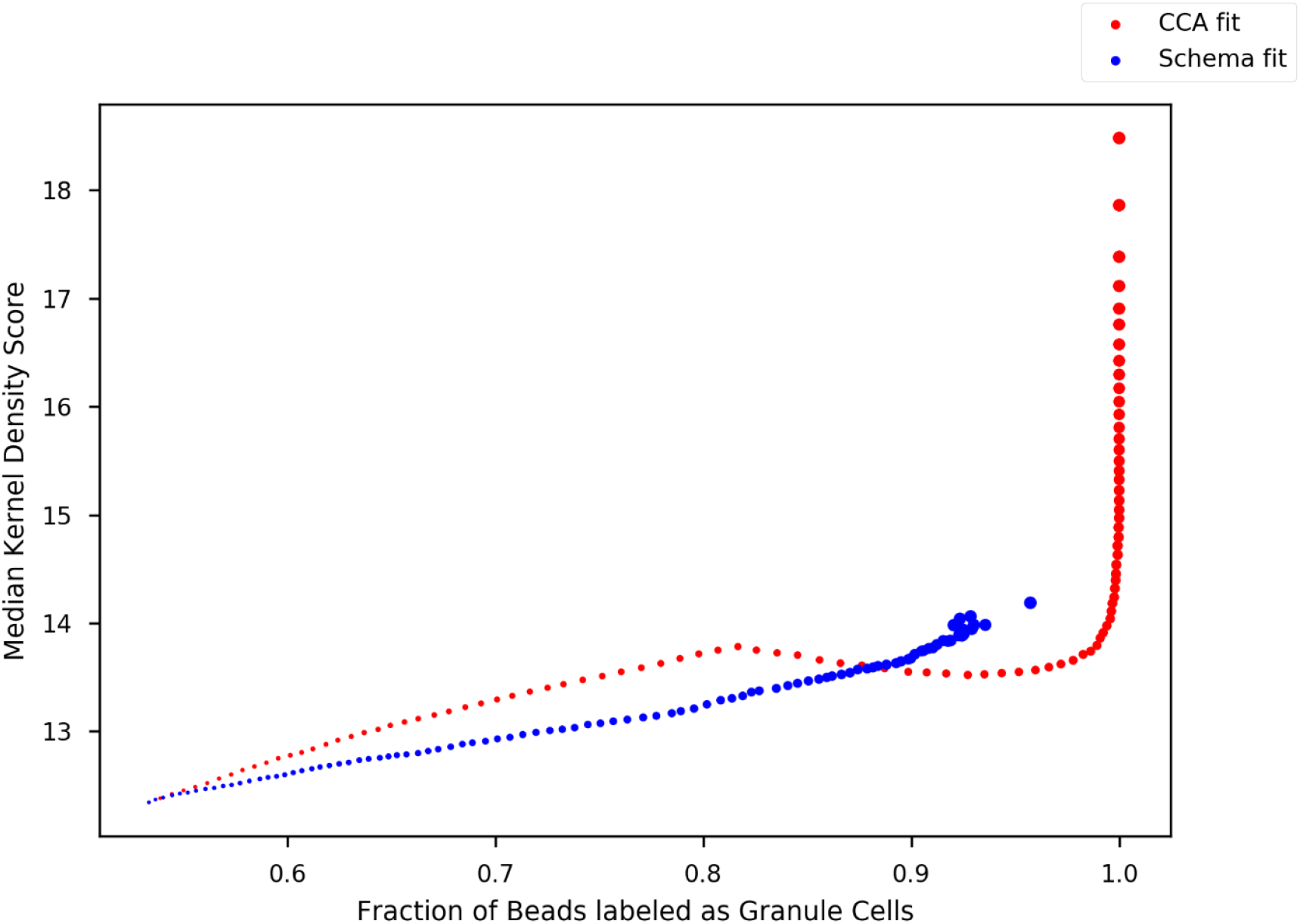
Comparison of Schema with CCA on Slide-seq data (Part 2) *Investigation of CCA and Schema cell loadings*: We sorted beads by their loading on the Schema-implied gene scores and investigated how the exposure to secondary modalities (cell-type labels and spatial density) varied in this ordering; we repeated the analysis for CCA cell loadings. For *k*=1,…,99, we selected cells with loadings in the percentile range [*k, 100*] and computed the frequency of granule-cell labels and the average Gaussian kernel density score of a cell in this set; higher values of these measures indicate stronger agreement with the cell type and spatial density modalities, respectively. In the plot, the size of a point is proportional to *k*. For both Schema and CCA, the higher cell loadings typically correlate with a higher granule-cell fraction and higher kernel density, as both the methods transform the primary gene-expression modality to align it with the secondary modalities. However, for Schema this relationship plateaus after a point because Schema’s regularization mechanism limits the distortion of the primary modality, constraining the extent of match with the secondary modalities. In contrast, the unconstrained framework of CCA produces loadings where the 99^th^ percentile cell loading has significantly higher spatial density exposure than the 95^th^ percentile. This may lead to overfitting as CCA computes gene rankings are overly determined by sample-specific artifacts. In contrast, the regularization mechanism of Schema produces gene rankings that are better preserved across samples.

**Supp Figure 9.**
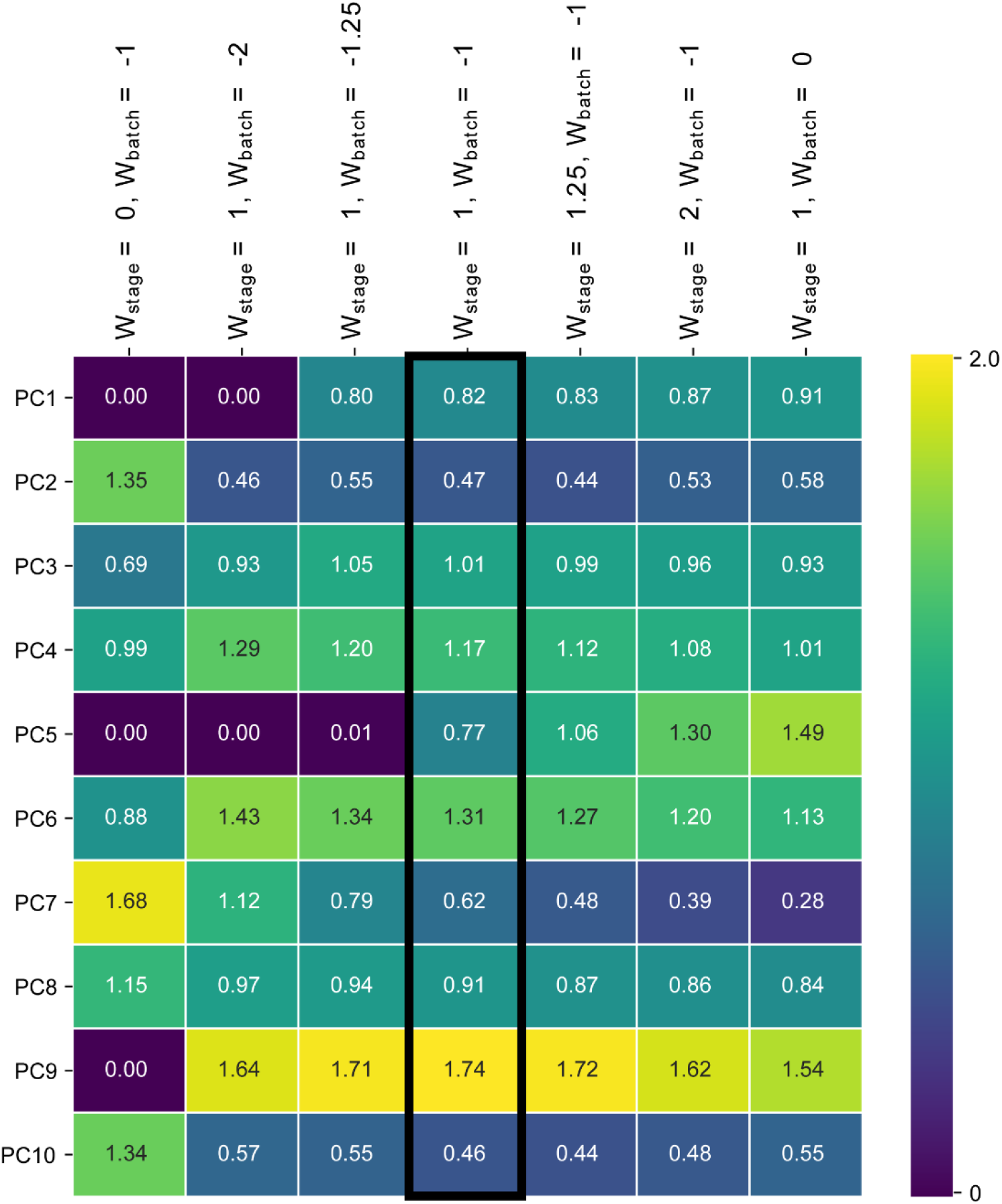
Differential expression analysis while accounting for batch effects and developmental stage: Schema feature-selection results for different weights of the developmental-stage and batch-effect modalities. The middle column is the one shown in the main text: equal (and opposite) weights for the batch-effects and developmental-stage modalities. The left-most column corresponds to using only the batch-effect modality while the right-most column corresponds to using only the developmental-stage modality.

**Supp Figure 10.**
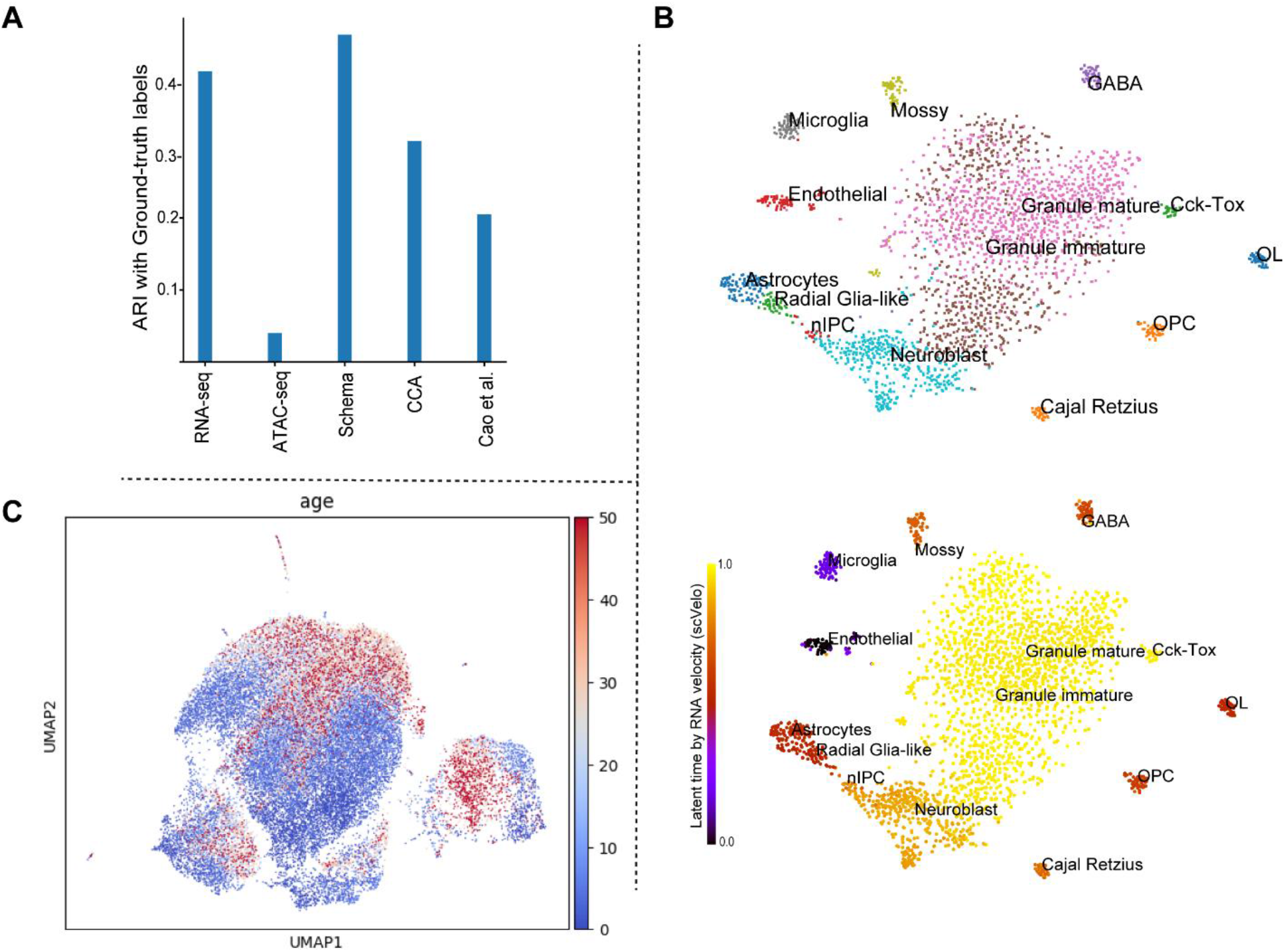
Evaluation of canonical correlation analysis (CCA) performance (also see Supp Figure 7, 8) (**A**) *Inferring cell types by synthesizing RNA-seq and ATAC-seq data*. The metric of evaluation here is the agreement between Leiden clustering on the synthesized dataset and ground truth cell-type labels, measured using the adjusted Rand index (ARI), with higher scores indicating greater agreement. This panel contains the same information as **Figure 2E** and is reproduced here for convenience. (**B**) *Inferring RNA velocity by synthesizing spliced and unspliced mRNA counts*. Spliced and unspliced data were correlated using CCA and the synthesized data was visualized with t-SNE. The bottom half of the panel colors cells by their scVelo latent-time estimate. This panel should be compared with **Figure 6B-F**, which display the corresponding plots produced by Schema synthesis of the data. CCA’s synthesis does not place cells with similar stages of differentiation as closely together as Schema. Spearman rank correlation between t-SNE distances and latent-time difference is 0.163 for CCA, while it is 0.397 when using just the spliced mRNA counts and 0.432 for the Schema transformation corresponding to a minimum correlation constraint of 0.95. (**C**) *Schema highlights secondary patterns while preserving primary structure*. RNA-seq data was synthesized with cell age metadata using CCA. Compared to a synthesis by Schema (**Figure 5B-D**), the CCA-based visualization less clearly communicates the developmental trajectory. We quantified the age-related structure in the transformed dataset by a diffusion pseudotime analysis. The Spearman rank correlation between the pseudotime estimate and the ground-truth cell age is only 0.059 for the CCA-synthesized data while it 0.365 in the original, untransformed dataset and 0.436 in the Schema-transformed dataset corresponding to a minimum correlation constraint of 0.99.

**Supp Table 3.**
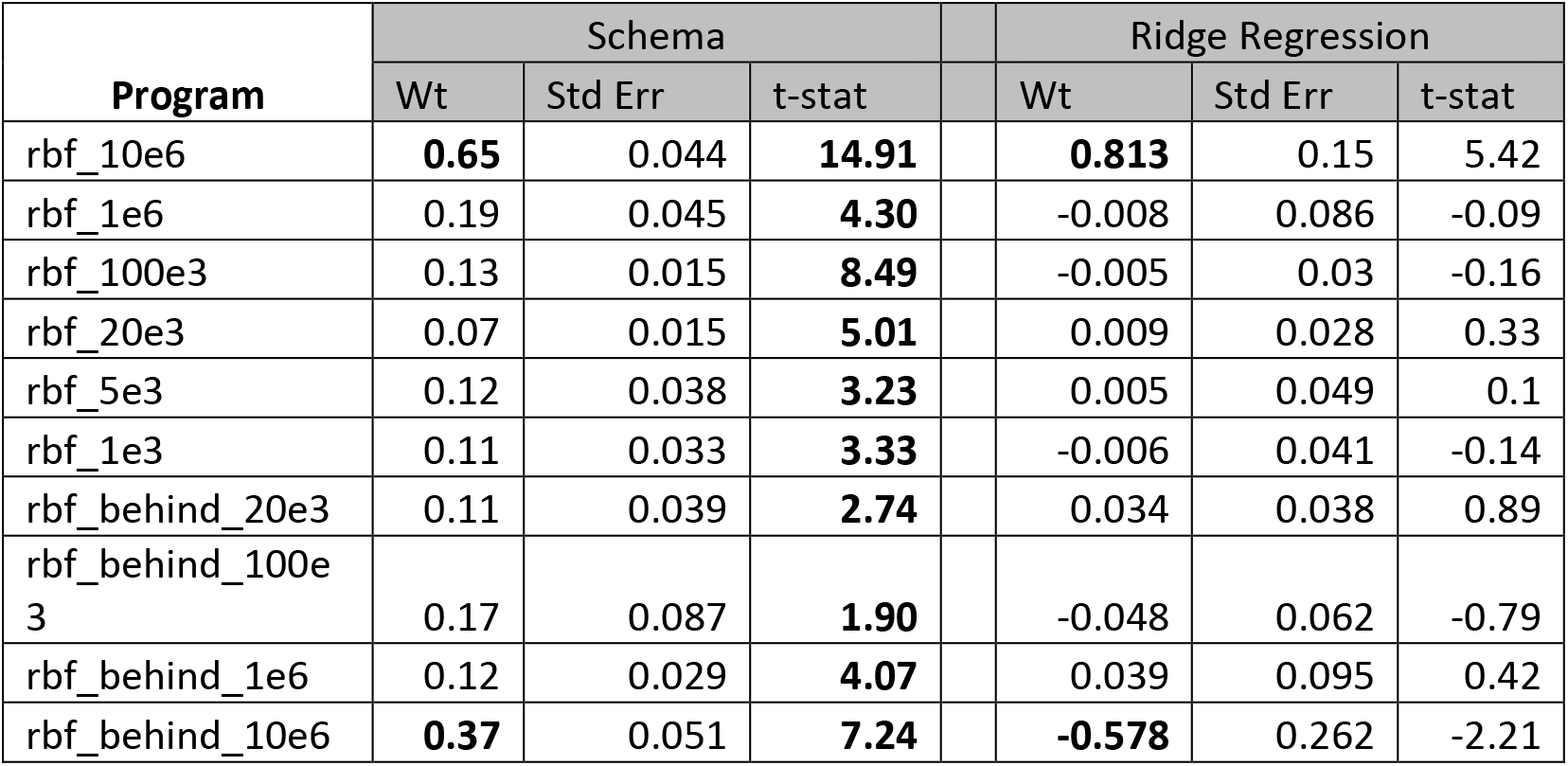
Comparison of feature weights estimated by Schema and ridge regression when integrating chromatin accessibility and gene expression data. Feature weights estimated by Schema are always non-negative, indicating just the feature’s importance in determining a gene’s expression variability. In regression, feature weights can be positive or negative, indicating whether the feature is relevant for high or low variability genes. Hence, when interpreting the ridge regression output we only considered the magnitude of a feature’s weight and not its sign. To compare the robustness of Schema and ridge regression estimates, we ran both these algorithms on subsets of genes bucketed by their strand orientation and chromosome. The regularization mechanism of Schema helps produce estimates that are more stable. The standard error and *t*-statistic (i.e., the ratio of *Wt* to *Std Err*) were computed from these.

| Program | Schema |  |  |  | Ridge Regression |  |  |
| --- | --- | --- | --- | --- | --- | --- | --- |
|  | Wt | Std Err | t-stat |  | Wt | Std Err | t-stat |
| rbf_10e6 | **0.65** | 0.044 | **14.91** |  | **0.813** | 0.15 | 5.42 |
| rbf_1e6 | 0.19 | 0.045 | **4.30** |  | -0.008 | 0.086 | -0.09 |
| rbf_100e3 | 0.13 | 0.015 | **8.49** |  | -0.005 | 0.03 | -0.16 |
| rbf_20e3 | 0.07 | 0.015 | **5.01** |  | 0.009 | 0.028 | 0.33 |
| rbf_5e3 | 0.12 | 0.038 | **3.23** |  | 0.005 | 0.049 | 0.1 |
| rbf_1e3 | 0.11 | 0.033 | **3.33** |  | -0.006 | 0.041 | -0.14 |
| rbf_behind_20e3 | 0.11 | 0.039 | **2.74** |  | 0.034 | 0.038 | 0.89 |
| rbf_behind_100e3 | 0.17 | 0.087 | **1.90** |  | -0.048 | 0.062 | -0.79 |
| rbf_behind_1e6 | 0.12 | 0.029 | **4.07** |  | 0.039 | 0.095 | 0.42 |
| rbf_behind_10e6 | **0.37** | 0.051 | **7.24** |  | **-0.578** | 0.262 | -2.21 |

### Supp Text 1

#### Applicability of existing metric learning algorithms

Unfortunately, existing metric learning methods ^22–25^ are not well suited to the challenge of synthesizing multi-modal single-cell data. Many of the considerations we discussed when comparing Schema to CCA also apply here: these methods, some of which list below, are designed to synthesize two datasets at a time, necessitating an *ad hoc* approach to integrating additional modalities. Like CCA, standard metric learning approaches do not limit the distortion of the primary modality. Setting a researcher-specified limit on this distortion is an important regularization mechanism in Schema, increasing the robustness of its results and ensuring that insights from the primary modality are not lost.

We designed Schema so it could scale to the large and ever-growing single-cell datasets. Towards that end, Schema deviates from existing metric learning approaches in computing a scaling transform and not a general affine transform. While affine transforms potentially offer more general alignment, Schema’s ability to accept arbitrary distance metrics on the secondary modalities partly compensates for this limitation on the primary modality transform. Additionally, the reach of a scaling transform is enhanced by featurizing the primary modality so that each feature represents a source of variance that is axis-aligned and orthogonal to others (e.g., using PCA or NMF). Most crucially for our needs, scaling transforms can be computed efficiently; they can be optimized using fast quadratic programming methods whereas an affine transform would need to be optimized using the much slower framework of semi-definite programming. Additionally, our choice of correlation as the measure of agreement allows for a sampling approach that further enhances scalability while producing provably accurate results (**Supp Text 3**).

These design choices allow Schema to scale up to large single-cell datasets. We ran a set of metric learning algorithms on one of the Slide-seq samples^10^ (puck 180430_1: 22943 cells x 18133 genes), using implementations made available on the creators’ websites or in the Python package *metric_learn*^47^. We tested the following methods: neighborhood component analysis^22^ (NCA); metric learning for kernel regression ^48^ (MLKR); local Fisher discriminant analysis^49^ (LFDA); large margin nearest neighbors^24^ (LMNN); and information theoretic metric learning^23^ (ITML). On a Linux server with 24 Intel Xeon 2.40 GHz cores and 386 GB RAM, each of these methods either crashed or failed to produce a meaningful output within 6 hours. In contrast, the aggregate runtime of an ensemble of Schema runs over different choices of the minimum correlation hyperparameter was 34 minutes on this dataset (Supp Table 4); in each run, Schema sampled a subset of the pairwise distances between points.

**Supp Table 4.** Runtime comparison of Schema with CCA, SpatialDE and Trendsceek. The values above were averaged over the three previously-described mouse cerebellum samples from the Slide-seq dataset. All programs were run on a Linux server with 24 Intel Xeon 2.40 GHz cores and 386 GB RAM. Each program was allowed to use as many cores as were available; Schema, CCA and SpatialDE did so, but Trendsceek did not. The server is a shared resource and while we did periodically check it to confirm that ample system resources were available, the runtime estimates above may be influenced by the load from other programs. For Schema, the runtime includes the time for pre-processing and encompasses the complete ensemble of sub-runs on different parameter choices. The runtime for CCA also encompasses the entire pipeline: pairwise modality combinations and then a final integration. For Schema and CCA, we were able to use all the data of each sample. For SpatialDE and Trendsceek, we experimented with small subsets of data and increased the subset size until the demand on the shared resource (server) became infeasible. The subset of cells for SpatialDE and Trendsceek were randomly chosen, with an equal split between granule and non-granule cells; for both, genes were selected based on high expression variability.

| **Program** | Average per sample |  |  |
| --- | --- | --- | --- |
|  | # of cells | # of genes | Run-time (in minutes) |
| Schema | 20823 | 17607 | 34 |
| CCA | 20823 | 17607 | 50 |
| SpatialDE | 16000 | 9000 | 244 |
| Trendsceek | 2000 | 3000 | 338 |

### Supp Text 2

#### Setting up the quadratic program

We introduce some notation to condense the expressions. Define *w*∈*R^k^* where 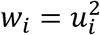, *δ_ij_* ∈ *R^k^* with 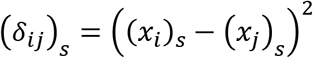 (i.e. the squared elements of *x_i_* – *x_j_*) and, for convenience, let *P* be the set of pairs of observations *P* = {{*i*, *j*} : 1 ≤ *i* ≤ *j* ≤ *N*}. Using the fact that the covariance between variables *X* and *Y* is given by *Cov*(*X*, *Y*) = *E*[*XY*] – *E*[*X*]*E*[*Y*], and the variance as *Var*(*X*) = *E*[*X*^2^] – *E*[*X*]^2^, we can expand:

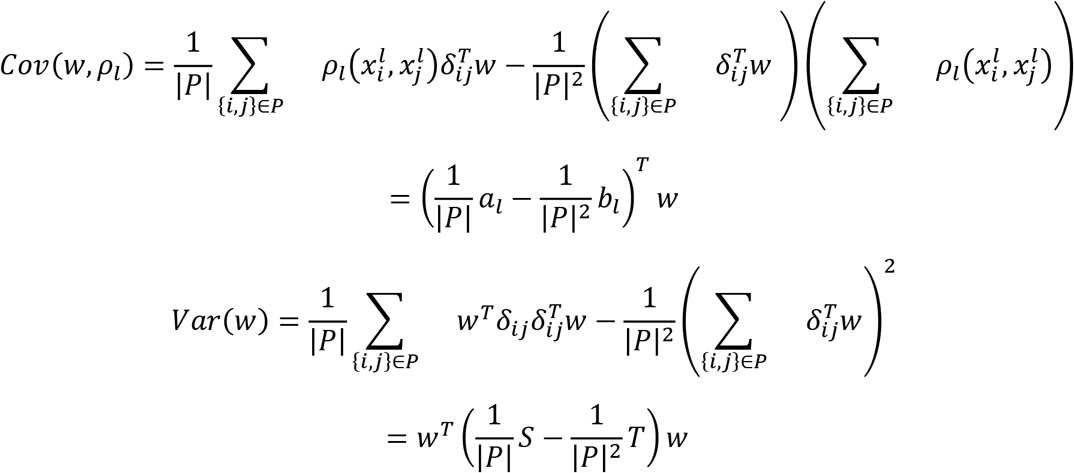

where *a_l_* and *b_l_* are *k*-dimensional vectors that depend only on *D_l_*; and *S* and *T* are *N* × *k* matrices that depend only on *D*_1_.

Explicitly, we derive:

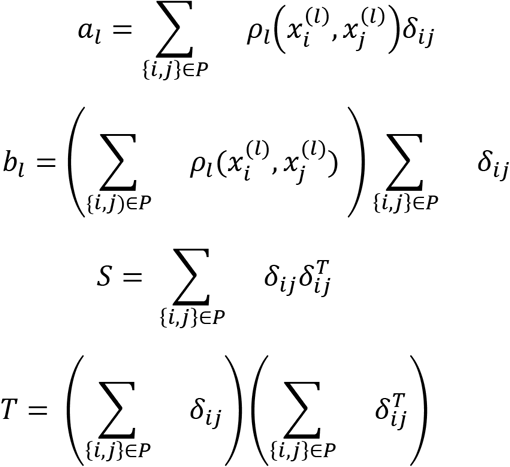

We recall the general optimization problem:

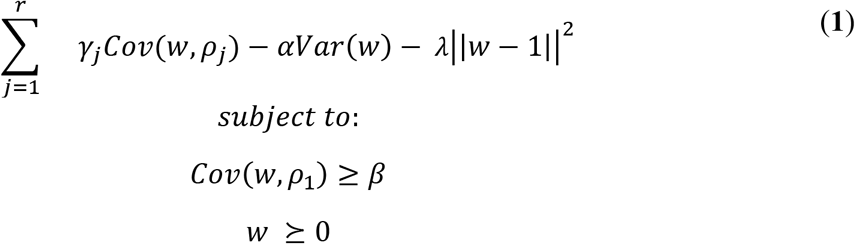

and the framework for quadratic programming that this needs to be mapped to:

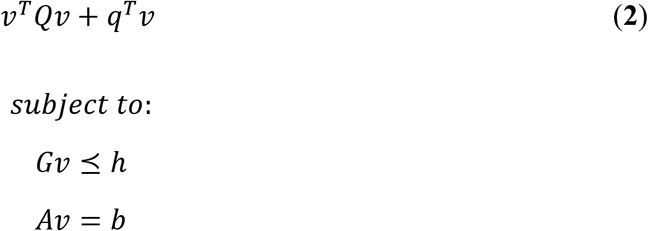

where *Q* is positive semidefinite.

The mapping from the general optimization problem (**1**) to the QP framework (**2**) is as follows:

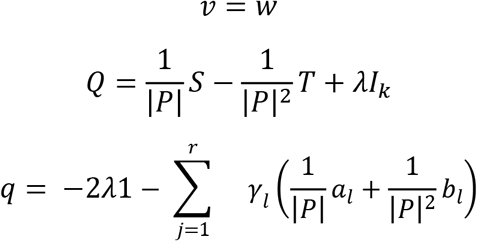

We also require that *Q* be positive semidefinite (psd). This is also straightforward to show. We can write:

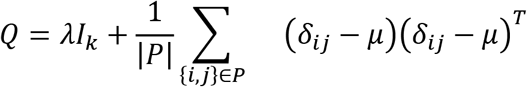

where 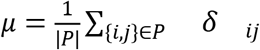, so it is a sum of psd matrices.

For the linear constraint, we express *G* as a block matrix:

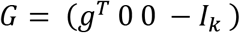

where

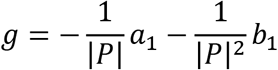

Lastly, we have, for the right side of the inequality constraint, each coordinate of ***h***:

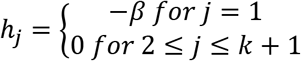

We have no equality constraints in our optimization, so *A* and *b* from (**2**) are not needed.

### Supp Text 3

#### Theoretical analysis of Schema’s scalability: concentration bounds

Our approach is to show that, given a 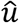 that has been calculated based on a random sample, the correlation coefficient between *all* pairwise distances cannot be too different than the correlation coefficient computed on the sample. To do this, we use Chernoff bounds, which bound how far away a random variable can be from its expectation, on the covariance and variance terms of correlation coefficient given. This gives us a bound on how far away the correlation coefficient on the whole population can be from the one calculated on the sample.

Let *P* be a random subset of all possible interactions. For now, we assume that interactions are chosen uniformly at random. Solving the optimization problem from the above section (Equation **(1)**) with our sample *P* yields 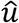, an estimator for the true optimal transform *u*. We show that 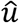 approximates *u* well by showing that the pairwise distances of 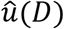 have high correlations with the secondary datasets as long as 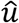 has high correlations on the subsample.

Formally, we will guarantee, for any *α, δ* > 0 and sample size at least 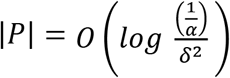.

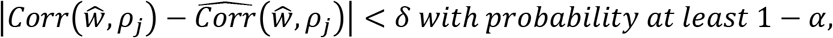

where 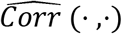 is the sample correlation coefficient (i.e. the Pearson correlation computed on the sample *P*).

This is a powerful result, made possible by our restriction to scaling transforms, which are easy to analyze. First of all, note that we only need a sample-size *logarithmic* in our desired confidence level in order to get strong concentration, allowing analysis of massive RNA-seq datasets.

To begin our analysis, let *W* be a *k* × *k* psd matrix (in our specific case it will be diagonal, but this analysis will generalize to any psd matrix, which motivates the generalization to all psd matrices in future work). We also assume randomly draw pairwise differences *δ* = *x_i_* – *x_j_*, choosing these *x_i_*, *x_j_* uniformly from the set of pairs of points in our primary dataset. Here, we focus on the correlation between the transformed dataset and the primary dataset. Analyses for correlations between the transformed data and the secondary datasets will be similar.

Consider the form of the (population) correlation:

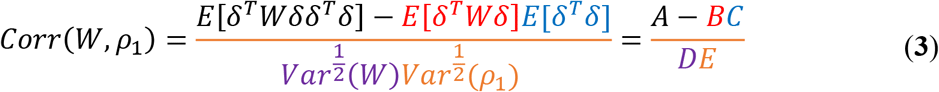

If, for our samples, we can determine confidence intervals of size 2*ϵ* for each of the terms *A, B, C, D, E* then we can bound the distance away from the correlation on the *entire* set of pairwise distances. This distance is maximized when *A* is as small as possible, and *B, C,D,E* are as large as possible. So, by expanding using Taylor expansions and removing terms of size *O*(*ϵ*^2^) or smaller, we get:

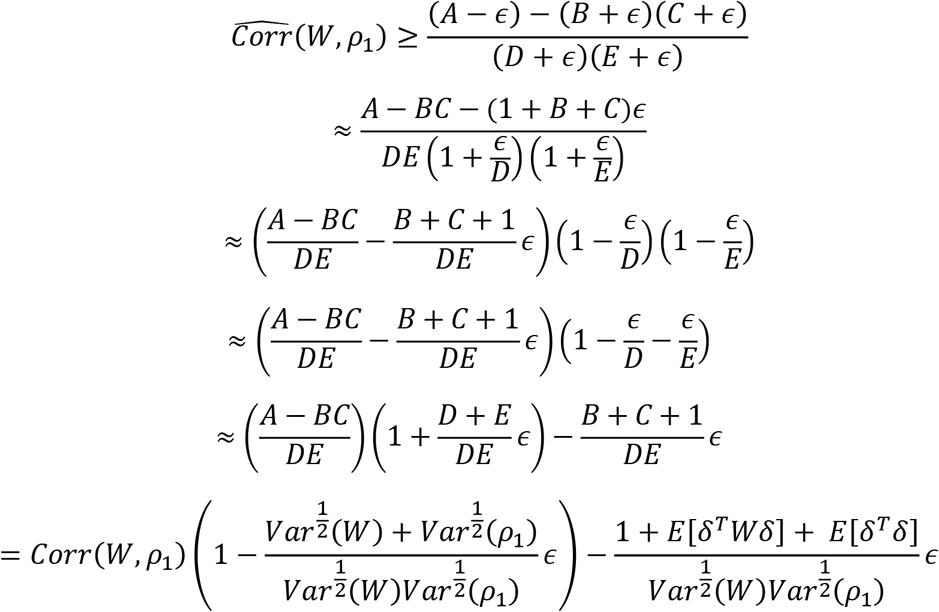

Thus, for a desired overall confidence level *η*, the relationship between *ϵ* and *η* is given by:

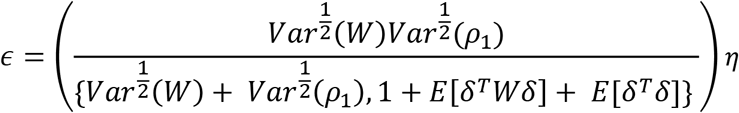

To show that we can bound each of the terms *A, B, C, D, E*$ we use *Hoeffding’s inequality* to limit how far away the terms can be from their expectations. Let *X*_1_ …, *X_n_* be i.i.d. random variables drawn from bounded range [*a*, *b*], and set *s* = *b* – *a*, and let 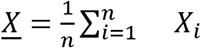. Then Hoeffding’s inequality states:

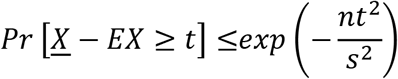

This can be converted into giving a (one-sided) confidence interval of length *t* by substituting the probability on the left with a desired confidence level *α*, and solving for *n*, which gives a statement:

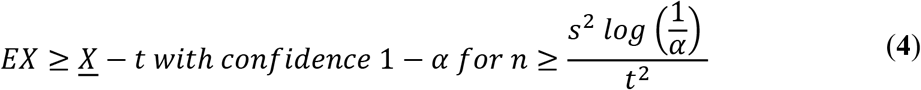

We begin by applying the inequality on term *A* = *E*[*δ^T^Wδδ^T^δ*] by bounding *δ^T^Wδδ^T^δ*. It is clear that 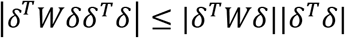, so we can bound each individually. Note that we can assume without loss of generality that *W* is diagonal here, because otherwise (since it is psd), we could write *W* = *UDU^T^*, where *D* is diagonal and *U* is unitary; setting *y* = *Uδ* yields 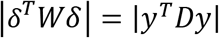, and, by unitarity, 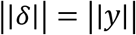.

Then, by Cauchy-Schwarz:

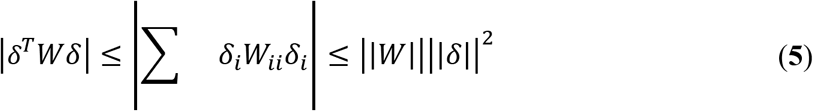

where 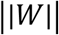 is the matrix-norm, i.e. 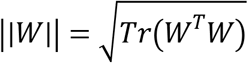. So for a diagonal matrix, 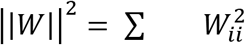. We can bound 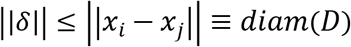.

Thus, 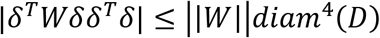.

To get a confidence interval of size *ϵ*, we plug into (**4**), so we require:

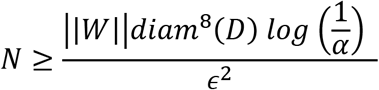

Note that the diameter is an *extremely* coarse bound for the above bound. Morally, one can replace “diameter” with “variance”, and the user has control over 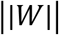 by choice of hyperparameters. We also note that the sample complexity improves drastically if we focus only on *local* distances, a future area of exploration.

The same analysis can be used for terms *B* and *C* in (**3**), but the dependency on the diameter is not as bad for those terms, so term *A* is the worst case.

Now, we consider the variance terms *D* and *E*. For term *E*, note:

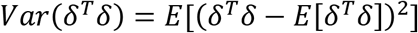

Again, |*δ^T^δ* – *E*[*δ^T^δ*]| is bounded by the maximum squared distance in the dataset *diam*^2^(*D*), so we can use the Hoeffding inequality from above in the same way.

And term *D* takes the same form as above, but with *δ^T^Wδ* instead of *δ^T^δ*. As noted in (**5**), this is a bounded random variable as well, but here with bound 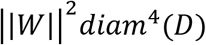.

Thus, in order to get a uniform confidence interval across all the terms, we require:

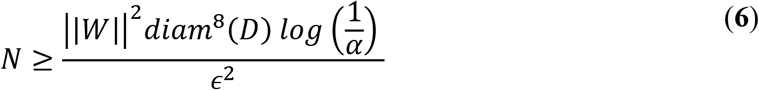

### Supp Text 4

#### Enriched functions and pathways for differentially expressed genes in dense granule cells

The densely-packed granule cell genes identified by Schema are strongly enriched for signal transmission, potentially indicating greater neurotransmission activity within these cells. In particular, REACTOME ^27^ pathway enrichment analysis (top 1000 genes, mapped to human) include vesicle-mediated transport (FDR *q* = 5.11 x 10^−4^), ion-channel transport (FDR *q* = 1.82 x 10^−3^), and cellular responses to external stimuli (FDR *q* = 6.44 x 10^−15^) (**Supp Table 1**, **Supp Figure 1**). An enrichment analysis of this gene set against the Gene Ontology (GO) database, performed using the GOrilla web-tool ^28^ and visualized using REViGO ^29^ also identified terms consistent with such activity: ion transport (GO:0022853, FDR *q* = 1.8 x 10^−17^), electron transfer (GO:009055, FDR *q* = 2.87 x 10^−11^) and enzyme binding (GO:0019899, FDR *q* = 2.72 x 10^−11^). (**Supp Table 2, Supp Figure 2**). Interestingly, we also observed enrichment in REACTOME pathways related to autophagy (FDR *q* = 3.19 x 10^−4^), ubiquitination (FDR *q* = 1.94 x 10^−4^) and protein metabolism (FDR *q* = 3.3 x 10^−7^). In particular, we observed enrichment for the process of Neddylation (FDR *q* = 2.26 x 10^−3^), shown to have a role in nuclear protein aggregation ^30,31^.

